# *HDAC5*-encoded Microprotein NISM Mediates Nucleolar Formation and Ribosomal RNA Synthesis

**DOI:** 10.64898/2026.02.21.707204

**Authors:** Kevin Cao, Dat Ha, Jesse Hulahan, Lilianna Houston, Gregory Tong, Jiayi Weng, Natasha Huey, Hannah E. DeMerit, Pedro Ortega, Rémi Buisson, Kingshuk Ghosh, Thomas F. Martinez

**Author notes:** Lead Contact: Thomas F. Martinez.

## Abstract

Ribosome biogenesis is the process by which ribosomal RNA (rRNA) and ribosomal proteins are synthesized, processed, and assembled into functional ribosomes. This process begins in the nucleolus, a multiphase liquid condensate. Here, we discover an arginine-rich disordered nucleolar microprotein encoded within the *HDAC5* 5′-UTR that we termed Nucleolar Integrity and Stress Microprotein (NISM). NISM overexpression leads to impaired rDNA transcription, triggering nucleolar stress, p53 activation, and suppressed proliferation. NISM knockout causes disruption of nucleolar structure and also induces p53 activation. Mechanistically, NISM interacts with the DExH-box RNA helicase DHX9 and regulates its activities related to pre-rRNA synthesis. Computational analyses and polymer physics-based mathematical modeling revealed that NISM coordinates nucleolar formation and pre-rRNA synthesis by enhancing the liquid-liquid phase separation of DHX9. This study establishes NISM as a regulator of nucleolar biology and deepens our understanding of how disordered microproteins can facilitate the formation of membraneless organelles.

## INTRODUCTION

Microproteins are a previously overlooked class of proteins encoded by small open reading frames (smORFs) less than 100 codons in length^1,2^. Microprotein-coding smORFs are found within regions of the transcriptome previously thought to be non-coding, including 5′- and 3′-untranslated regions (UTRs), long non-coding RNAs (lncRNAs), and overlapping CDS regions in alternative reading frames. Microproteins function in a variety of critical cellular processes, including DNA repair^3,4^, cellular respiration^5^, mitochondrial metabolism^6^, endoplasmic reticulum stress^7^, and RNA decay^8^. While >10,000 putative human microproteins have been identified to date^9^, the vast majority remain uncharacterized and difficult to study, in part because of their lack of homology to known proteins and protein domains. We and others have found that mammalian microproteins are enriched in alanine, glycine, arginine, and proline^10,11^ and are most often predicted to be disordered^12,13^. Intrinsically disordered regions (IDRs) within larger annotated proteins are known to be critical regulators of cellular functions, such as transcriptional regulation, cell signaling, molecular complex formation, nucleolar organization, and more^14^. Their lack of structure enables IDRs to engage in multivalent, tunable, and malleable interactions that make them ideal for responsive regulation that would be more difficult for folded domains. Additionally, IDR-mediated multivalent interactions help promote phase separation of membraneless organelles (MLOs), or biomolecular condensates, which are now known to be critical for myriad biological processes. Based on our understanding of IDRs within annotated proteins, we rationalized that highly disordered microproteins are likely to act in similar fashions which can be inferred from their sequence properties.

Arginine-rich motifs (ARMs) within disordered regions are frequently found in RNA binding proteins and other proteins involved in RNA metabolism^15^. For instance, RGG/RG motifs are commonly found in the RNA binding domains and repeats of the motif can mediate interactions that promote phase separation^16^. Similarly, RS repeats can also bind RNA and are found in pre-mRNA splicing related proteins. K/R-rich basic patches also occur often in RNA related proteins, such as the HIV-1 Tat and Rev proteins, which have roles in transcriptional activation and RNA export, respectively^15^. Considering the frequency of arginine and glycine residues in microproteins and their propensity for disorder, we hypothesized that many are likely involved in regulating aspects of RNA metabolism.

In this study, we searched for ARM-containing disordered microproteins with roles in RNA metabolism. This led us to the discovery a microprotein encoded within the 5′-UTR of *HDAC5*, which we termed Nucleolar Integrity and Stress Microprotein (NISM). NISM was found to localize to the nucleolus and interact with the DExH-box RNA helicase DHX9, which has roles in resolving R-loops, three-stranded DNA:RNA hybrids, and G-quadruplexes as well as other activities^17–19^. Perturbation of NISM revealed that it regulates nucleolar dynamics, with its loss causing disruption of nucleolar structure and its overexpression leading to impaired nucleolar R-loop homeostasis and inhibition of 47S pre-rRNA synthesis. Computational analyses and polymer physics-based theoretical calculations indicated that NISM binding increases DHX9’s propensity for phase separation, providing a basis for these observed effects. Our findings provide what we believe is the first evidence of a microprotein that modulates the phase separation of another protein and suggest that more microprotein regulators of RNA metabolism are likely hidden.

## RESULTS

### Identifying microproteins with arginine-rich motifs

To identify arginine-rich microproteins which may have roles in RNA metabolism, we first updated our previously developed in-house microprotein database^10^. We recently reported that different bioinformatics tools for ribosome profiling (Ribo-seq) analysis can report vastly different sets of translated smORFs from the same dataset, and that smORFs called translated by multiple tools tend to have higher confidence predictions^20^. Therefore, we reanalyzed our Ribo-seq datasets collected from HEK293T, HeLa-S3, and K562 cells using both RibORF 1.0 and RiboCode. Our updated analyses resulted in 10,587 smORFs called translated by both tools with ∼51% being detected in multiple Ribo-seq datasets (**Supplemental Data 1, Figure S1A**). We next filtered our database for microproteins that contain ARMs, specifically two or more RG, RS, or RRR motifs as well as those with R representing ≥20% of the sequence. In total, 2,497 microproteins (∼24%) contain at least one ARM, 741 microproteins (∼7%) possess multiple motifs, and 20 microproteins (∼0.2%) contain all 4 motifs (**Supplemental Data 1, Figure S1B**). These analyses enabled us to curate a list of high-confidence arginine-rich microproteins with potential roles in RNA metabolism to investigate further. Moreover, users can analyze these microproteins for additional predicted features, such as IDRs^21^ and subcellular localization^22^ using computational tools.

### *HDAC5* 5′-UTR encodes an arginine-rich disordered nucleolar microprotein

Our search for arginine-rich microproteins led us to NISM, a 36 amino acid microprotein encoded by a smORF within the 5′-UTR of *HDAC5*. NISM was chosen for further investigation for multiple reasons. First, its smORF was called translated in all three analyzed cell lines (**Supplemental Data 1**) and it is also included in the Phase I Consensus set of translated smORFs curated by the TransCODE Consortium^9,23^. Notably, close inspection of the genomic locus for the smORF revealed an additional upstream in-frame AUG codon, which would result in a 53 codon smORF. Ribo-seq evidence in HEK293T cells shows most of the read coverage beginning at the downstream AUG, and cells pre-treated with the translation initiation inhibitor harringtonine also show read coverage predominantly at this same AUG (**Figures 1A and 1B**). However, there is a small amount of read coverage around the upstream AUG in both the Ribo-seq sample and the harringtonine sample. A similarly small amount of read coverage near the upstream AUG codon can also be observed in the Ribo-seq A-site plots from U2OS and K562 cells (**Figure S2A**). Given the possibility of translation initiation from the upstream AUG, both the potential short and long isoforms were investigated in this study. Further supporting the likelihood of NISM being a functional microprotein, its amino acid sequence is well conserved across distant mammalian species, including mouse, dolphin, and blue whale (**Figure 1C**).

**Figure 1.**
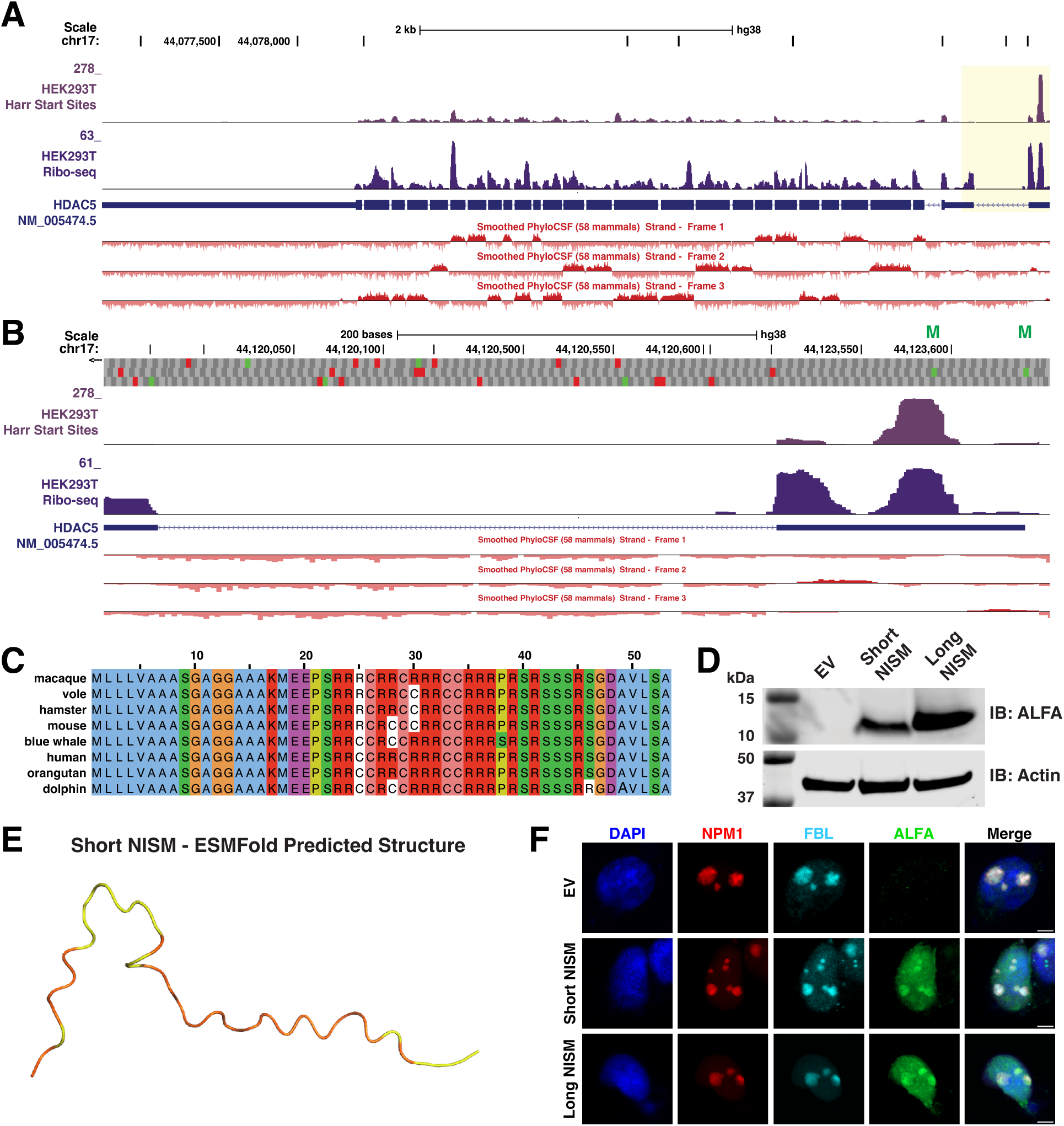
NISM is an intrinsically disordered nucleolar microprotein encoded on the 5′-UTR of *HDAC5*. **(A-B)** BedGraph tracks of HEK293T Ribo-seq coverage across the entire HDAC5 transcript (A) and zoomed in on the NISM smORF within the 5′-UTR (B). In (A), the NISM smORF is highlighted in yellow. *HDAC5* is on the negative strand and thus the 5′ to 3′ orientation runs right to left. The top tracks show read coverage for cells pre-treated with harringtonine (Harr) to capture translation initiation sites and the bottom tracks show read coverage for untreated cells lysed in the presence of cycloheximide to capture elongating ribosomes. PhyloCSF scores for each reading frame are shown below the RefSeq transcript tracks. Possible start codons in the NISM smORF are denoted by ‘M’ in (B). **(C)** Sequence alignment showing amino acid level conservation of NISM across distant mammals. **(D)** Immunoblot analysis of short and long NISM-ALFA expression in HEK293T cells. **(E)** ESMFold predicted structure of short NISM. Low confidence predictions are colored yellow and very low confidence predictions are colored orange. **(F)** Representative immunofluorescence images of NPM1 (red), FBL (cyan), and NISM-ALFA (green) in HEK293T cells transfected with empty vector (EV), short NISM-ALFA, or long NISM-ALFA for 48 h. Nuclei were counter-stained with DAPI (blue). Scale bar, 5 µm. All data are representative of at least two biological replicates.

Consistent with this observation, the mean PhyloCSF score^24^ for the short smORF isoform is 0.241, representing a modest likelihood for being a protein-coding ORF (**Supplemental Data 1**). Finally, exogenous expression of both short and long NISM C-terminally fused to ALFA (NISM-ALFA), a charge neutral alpha-helical epitope tag^25^, resulted in robust detection by immunoblot in HEK293T cells (**Figure 1D**). These data suggested that NISM is a functional arginine-rich microprotein.

As mentioned above, ARMs within disordered regions of proteins are frequently involved in RNA metabolism. NISM contains an arginine-rich low complexity region^26^, representing ∼36% of the short isoform sequence and ∼25% of the long isoform, which we hypothesized to be structurally disordered. Supporting this hypothesis, AIUPred^21^ predicted that both short and long NISM are entirely disordered, with scores well above the 0.5 threshold for disordered regions across the length of the microprotein (**Figure S2B**). Similarly, ESMFold^27,28^ predicted no structural features in either NISM isoform (**Figures 1E and S2C**).

Next, to begin uncovering its possible role in RNA metabolism, we used immunofluorescence to determine NISM’s subcellular localization. In HEK293T cells, NISM-ALFA was found to localize to the nucleus with significant enrichment in the nucleolus, which was identified by staining of nucleophosmin (NPM1) and fibrillarin (FBL) (**Figure 1F**). This observation is consistent with previous studies which showed that positively charged peptides with poly-arginine motifs are sufficient to drive proteins to the nucleolus^29–31^. Furthermore, the nucleolar localization sequence detector tool (NoD) also predicted that the arginine-rich regions in NISM are likely nucleolar localization sequences^32^ (**Figure S2D**). Lastly, to confirm that nucleolar localization was not an artifact of the ALFA tag, we found that expression of NISM fused to a C-terminal HA tag also localized to the nucleolus (**Figure S2E**). Altogether, these data reveal that NISM is an arginine-rich disordered nucleolar microprotein.

### Overexpression of NISM induces canonical nucleolar stress

Given that NISM-ALFA showed prominent localization to the nucleolus, we hypothesized that NISM has a role in ribosome biogenesis and that perturbation of its expression might induce nucleolar stress. Nucleolar stress is caused by disruption of any step during ribosome biogenesis, including rDNA transcription, rRNA maturation, and pre-ribosome assembly^33^. For example, nucleolar stress is frequently studied by low dose treatment with actinomycin D (ActD), which inhibits RNA polymerase I transcription of rRNA. In response to this stress, nucleolar proteins become redistributed into the nucleoplasm to inhibit MDM2, allowing for p53 stabilization and downstream cell cycle check point activation to attempt resolution of the stress^34^.

We investigated the effects of NISM perturbation in U2OS osteosarcoma cells, as nucleolar stress has been studied extensively in this model and these cells express wild-type p53^30,33,35–37^. Transient overexpression of either the short or long form of NISM-ALFA in U2OS cells resulted in significantly smaller and rounded nucleoli, a hallmark of canonical nucleolar stress (**Figures 2A and 2B**)^38^. We also noted the presence of nucleolar stress caps marked by concentrated FBL staining on the outer boundaries of nucleoli in response to ActD treatment. These caps are indicative of transcriptional inhibition^33^ and appear to occur in NISM-ALFA overexpressing cell as well, although less prominently.

**Figure 2.**
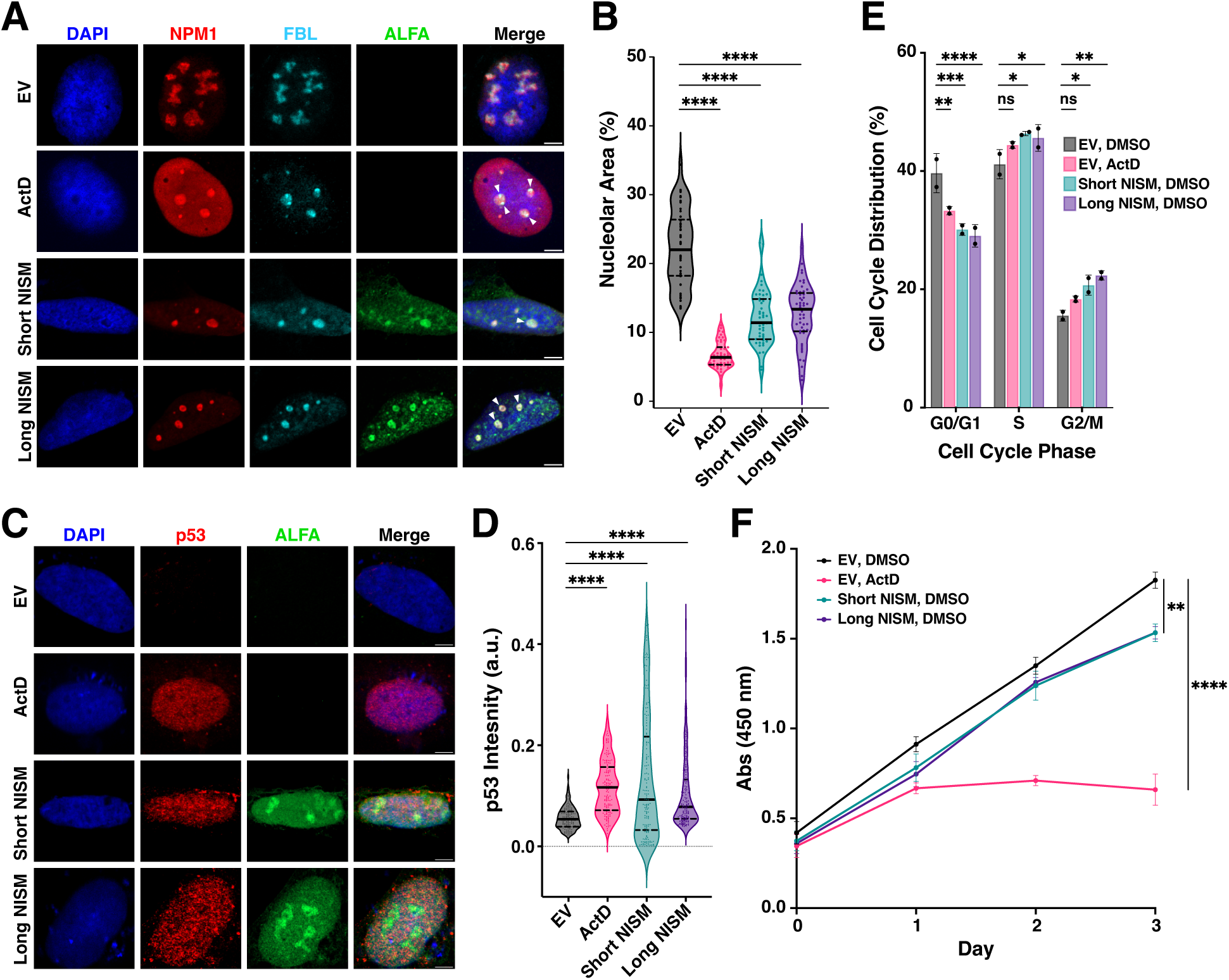
NISM-ALFA overexpression induces nucleolar stress and suppresses proliferation in U2OS cells. **(A)** Representative immunofluorescence images of NPM1 (red), FBL (cyan), and NISM-ALFA (green) in U2OS cells treated with 10 nM ActD or DMSO for 3 h and transfected with empty vector (EV), short NISM-ALFA, or long NISM-ALFA for 48 h. Nuclei were counter-stained with DAPI (blue). White triangles mark nucleolar stress caps. Scale bar, 5 µm. **(B)** Quantification of nucleolar area for cells in (A) as a percentage of total nuclear area represented by violin plots. Statistical comparisons were calculated by Mann Whitney test; n = 50 cells per condition. **(C)** Representative immunofluorescence images of p53 (red), and NISM-ALFA (green) in U2OS cells treated as in (A) for 72 h. **(D)** Quantification of p53 fluorescence intensity for cells in (C) represented by violin plots. Statistical comparisons were calculated by Mann Whitney test; n = 140-150 cells per condition. **(E)** Cell cycle analysis of propidium iodide-stained cells treated as in (A) represented by bar plots displaying the mean ± SD. Statistical comparisons were calculated by two-tailed Student’s t test; n = 2. **(F)** Cell proliferation analysis by WST-1 assay for U2OS cells treated as in (A). Asterisks indicate: *p < 0.05, **p < 0.01, ***p < 0.001, ****p < 0.0001, ns = not significant. All data are representative of at least two biological replicates.

We next sought to determine whether the nucleolar stress response pathway was activated in response to NISM-ALFA overexpression. First, we assessed U2OS cells for p53 stabilization. Immunofluorescence of p53 showed significant stabilization induced by both short and long NISM-ALFA overexpression, similar to ActD treatment (**Figures 2C and 2D**). Activation of p53 during nucleolar stress has previously been shown to induce accumulation of cells in the G2/M phase^39–41^. Consistent with this, NISM-ALFA overexpression caused an accumulation of cells in G2/M, a drop in G0/G1, and a modest increase in S-phase (**Figures 2E and S2F**). Following these results, we also observed moderate inhibition of U2OS proliferation upon microprotein overexpression (**Figure 2F**). Overall, these results supported disruption of ribosome biogenesis and subsequent triggering of the nucleolar stress response by NISM-ALFA overexpression.

### Knockout of *NISM* disrupts nucleolar structure and induces an alternative form of stress

Having established that exogenous overexpression of NISM induces nucleolar stress, we next investigated the effects of its knockout on nucleolar biology. CRISPR-Cas9 was used to create two clonal U2OS cell lines with identical homozygous 14 bp deletions within the NISM smORF (**Figures S3A-D**). This deletion occurred solely within exon 1 and caused only a slight drop in *HDAC5* transcript levels by qRT-PCR and no observable change in HDAC5 protein levels immunoblot (**Figures S3E and S3F**). As overexpression of the microprotein caused nucleolar shrinkage and activated the downstream stress response, we hypothesized that its knockout might also disrupt ribosome biogenesis and induce stress. Surprisingly, immunofluorescence of NPM1 and FBL showed significantly altered structure with more diffuse NPM1 and less compacted FBL staining as well as fewer defined nucleolar regions in both NISM knockout (NISM KO) cell lines (**Figures 3A and 3B**). A similar change in nucleolar structure and diffusion of granular component proteins has been reported previously by Potapova et al. upon treatment of U2OS cells with the CDK9 inhibitor AZD4573^38^. This effect was termed “bare rDNA scaffold,” as RNA polymerase I dissociates from rDNA. In our experiments, we found that the NISM KO cells appeared to have a milder effect, given that NPM1 staining was more diffuse but not entirely disrupted as under CDK9 inhibition (**Figures 3A and 3B**).

**Figure 3.**
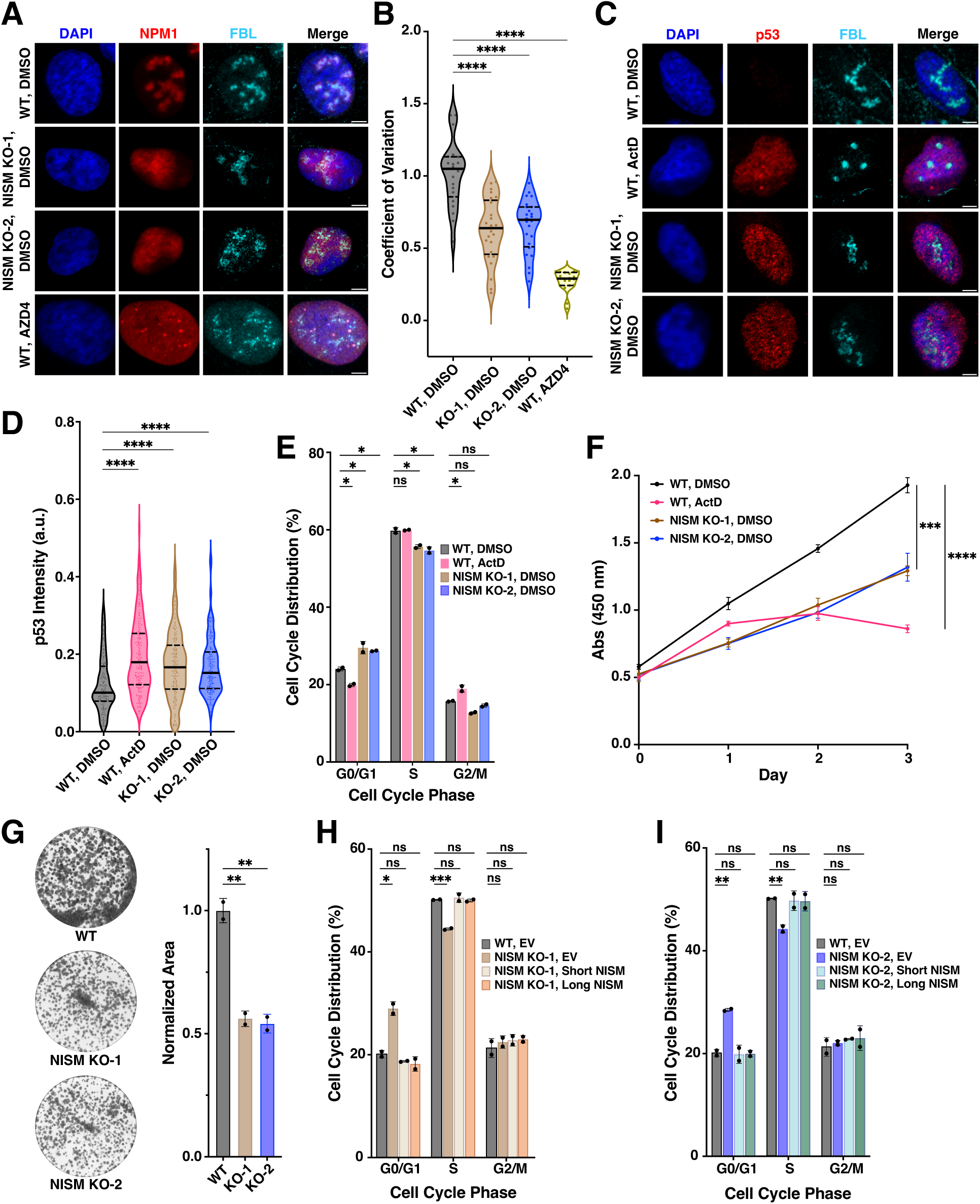
NISM knockout disrupts nucleolar structure and activates downstream stress pathway. **(A)** Representative immunofluorescence images of NPM1 (red) and FBL (cyan) in wild-type (WT) or clonal NISM knockout (NISM KO) U2OS cells treated with 5 µM of AZD4573 or DMSO for 5 h. Nuclei were counter-stained with DAPI (blue). Scale bar, 5 µm. **(B)** Quantification of the coefficient of variation of NPM1 area for cells in (A) represented by violin plots. Statistical comparisons were calculated by Mann Whitney test; n = 20-25 cells per condition. **(C)** Representative immunofluorescence images of p53 (red), and NISM-ALFA (green) in WT or NISM KO cells treated with 10 nM ActD or DMSO for 3 h. **(D)** Quantification of p53 fluorescence intensity for cells in (C) represented by violin plots. Statistical comparisons were calculated by Mann Whitney test; n = 150-175 cells per condition. **(E)** Cell cycle analysis of propidium iodide (PI)-stained cells treated as in (C) represented by bar plots displaying the mean ± SD. Statistical comparisons were calculated by two-tailed Student’s t test; n = 2. **(F)** Cell proliferation analysis by WST-1 assay for cells treated as in (C). **(G)** Colony formation assay of WT and NISM KO cells represented by bar plots displaying the mean normalized area covered by cells ± SD. Statistical comparisons were calculated by two-tailed Student’s t test; n = 3. **(H-I)** Cell cycle analysis of PI-stained WT and NISM KO cells overexpressing empty vector or short or long NISM-ALFA. Asterisks indicate: *p < 0.05, **p < 0.01, ***p < 0.001, ****p < 0.0001, ns = not significant. All data are representative of at least two biological replicates.

We next determined whether NISM KO cells had an active stress response to the disrupted nucleoli. As in NISM-ALFA overexpressing cells, we observed a significant and robust increase in p53 stabilization in the knockout cells (**Figures 3C and 3D**). However, in contrast with the overexpressing cells, NISM KO cells showed a significant increase in G0/G1 phase cells and a drop in S phase cells (**Figure 3E and S3G**). This change in cell cycle progression also resulted in a more severe reduction in cell proliferation than observed the NISM-ALFA overexpressing cells, as determined by both WST-1 and colony formation assays (**Figure 3F and 3G**). To validate the role of NISM in mediating these phenotypes, we reintroduced NISM-ALFA by exogenous expression and observed that both isoforms were able to fully rescue the cell cycle distribution effects of the knockout (**Figures 3H-I and S3H**). These results showed that NISM KO drastically alters the nucleolar structure and induces a different form of cell stress that causes severe suppression of cell proliferation.

### NISM Interacts with the DExH-box RNA Helicase Family Member DHX9

Microproteins frequently exert their functions by regulating the activities of larger proteins and protein complexes^2,42^. Thus, we conducted immunoprecipitation-mass spectrometry (IP-MS) experiments to identify potential binding partners and help uncover NISM’s role in the nucleolus. To help enrich for likely interactors, we first isolated nuclear lysates from U2OS cells overexpressing either short or long NISM-ALFA (**Figure S4A**). The microprotein was then pulled down using an anti-ALFA nanobody and co-eluted proteins were identified by mass spectrometry (**Figure S4B**). Significantly enriched proteins in the NISM-ALFA IP samples relative to eluate from empty vector transfected control samples were determined using SAINTexpress^43^. In total, 62 prey proteins were enriched with short NISM, 55 were enriched with the long isoform, and 49 were found in common for both NISM isoforms (**Figure 4A and Supplemental Data 2**). The large overlap in enriched proteins suggested that the additional N-terminal sequence in the long isoform is not a significant contributor to the microprotein’s interactome. Among the 49 common proteins were many ribosomal proteins, multiple HNRNP family members and additional RNA splicing proteins, and multiple DExD/H-box RNA helicase family members (**Figure 4B**). Given NISM’s localization to the nucleolus, we filtered the putative interacting proteins for those annotated as localized to the nucleolus in the Human Protein Atlas^44^ (**Figure S4C**). This limited our set to multiple regulators of rRNA synthesis, UBTF^45^, TOP1^46^, and PARP1^47^, ribosomal proteins, which are essential to pre-ribosome assembly in the nucleolus, and DExD/H-box helicases DDX5 and DHX9. Given that DHX9 and DDX5 were among the highest average spectral count hits in both the short and long isoform IPs and that both DExD/H-box helicase family members have reported roles in rRNA synthesis^48,49^, we focused on these hits to validate as potential direct interactors. We were able to validate the enrichment of DHX9 in the NISM-ALFA IP samples by immunoblot, but not DDX5 (**Figures 4C and S4D**). Further confirming the interaction, we found that reciprocal IP of endogenous DHX9 was also able to enrich NISM (**Figure 4D**).

**Figure 4.**
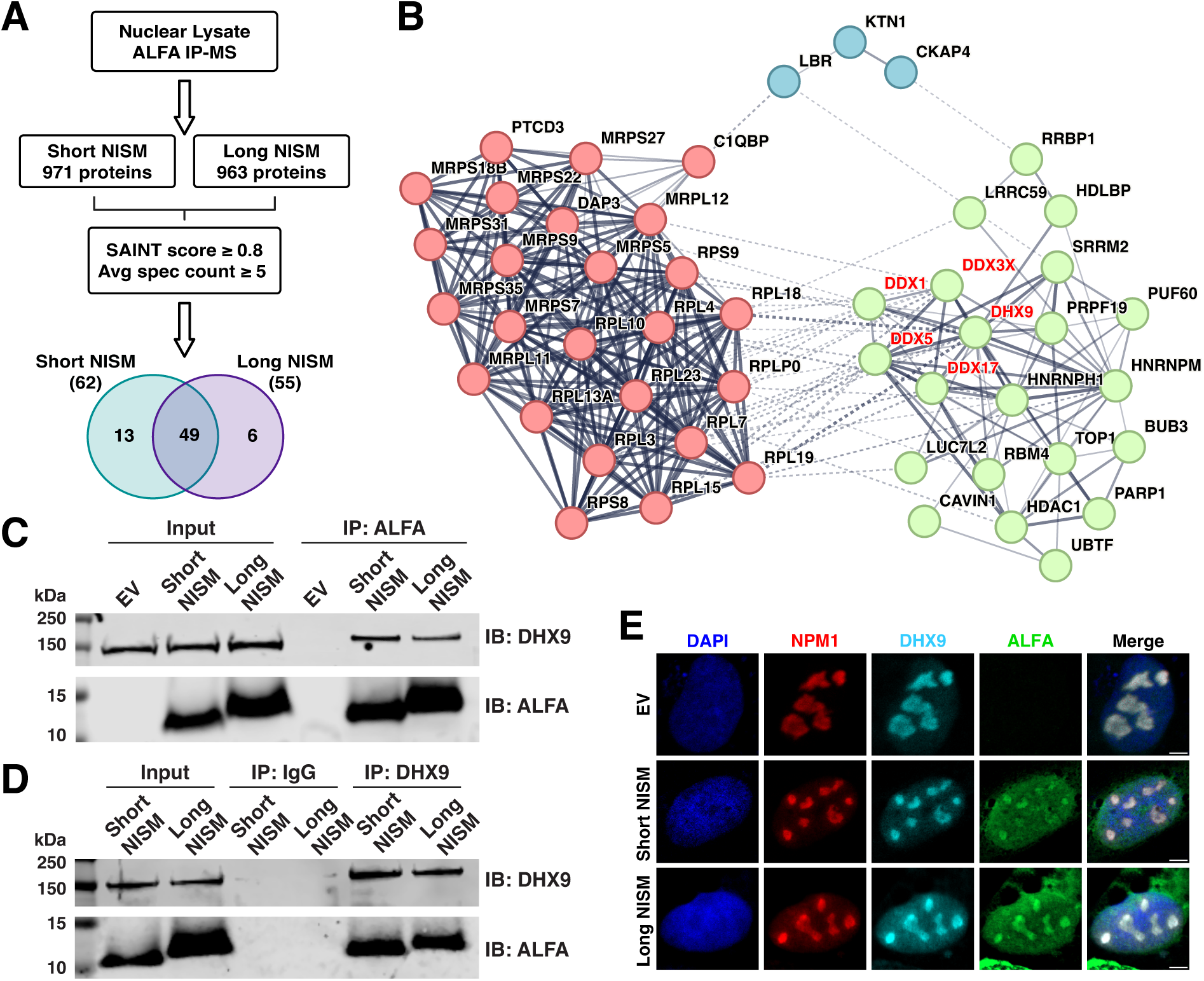
NISM interacts with DHX9. **(A)** Depiction of nuclear lysate immunoprecipitation-mass spectrometry (IP-MS) analysis workflow and Venn diagram showing the overlap in prey proteins significantly enriched by short and long NISM-ALFA; n = 3. **(B)** STRING database plot showing interactions between the 49 prey proteins pulled down by both short and long NISM-ALFA. DExD/H-box RNA helicase family members are highlighted in red and prey proteins were grouped by k means clustering. **(C-D)** Immunoblot validation of DHX9 enrichment in NISM-ALFA IP samples (C) and NISM-ALFA enrichment in DHX9 IP samples (D). **(E)** Representative immunofluorescence images of NPM1 (red), DHX9 (cyan), and NISM-ALFA (green) in U2OS cells overexpressing short or long NISM-ALFA. All immunoblots and images are representative of at least two biological replicates.

As immunoprecipitation was performed in homogenized nuclear lysates, we wanted to verify that NISM-ALFA and DHX9 co-localize within the nucleolus. We observed clear co-localization of both short and long NISM with DHX9 within nucleoli, as well as the nucleoplasm to a lesser extent, by immunofluorescence (**Figure 4E**). These results are consistent with published studies demonstrating that DHX9 is present in both the nucleoplasm^50,51^ and nucleolus^19,52^ in multiple cell types. The nucleolus itself contains additional sub-compartments within its membraneless structure, including the fibrillar center (FC), the surrounding dense fibrillar component (DFC), and the outer granular component (GC). Therefore, we wanted to look more closely within the nucleolus. A recent study used super-resolution microscopy to show nucleolar localization of DHX9 specifically to the periphery of the DFC (PDFC) in HeLa cells^53^. Our assessment of NISM-ALFA localization relative to nucleolar sub-compartment markers revealed that it overlaps with the DFC and GC (**Figure S4E and S4F**), which would allow for potential interaction with DHX9 in the PDFC. Altogether, these data support DHX9 as a key interactor of NISM in the nucleolus.

### Computational models support a direct interaction between DHX9 and NISM

While the IP and imaging experiments confirm there is an interaction between NISM and DHX9, they cannot determine whether the interaction occurs directly between the two proteins or whether there are other proteins or biomolecules mediating the interaction. To identify the key residues within short NISM that are necessary for its interaction with DHX9, we first attempted to use alanine scanning mutagenesis. By dividing short NISM roughly into thirds and mutating 11 to 13 sequential amino acids to alanine, we found that large mutations were only tolerated at the C-terminus from residues 24–36. Alanine mutations within the first two thirds of the sequence significantly reduced the stability of the microprotein (**Figure S5A**). We next tested shorter stretches of alanine mutations within NISM’s first 23 amino acids. Even by limiting the mutations to only 5 to 6 sequential residues, the NISM mutants were still unstable in cells (**Figure S5B**). The first 23 amino acids of short NISM encompass the arginine-rich low-complexity region, suggesting that these residues are critical for its stability and likely function. In addition, to understand how flexible the microprotein is along its sequence, we used STARLING^13^ to predict the conformational ensembles of short and long NISM (**Supplemental Videos 1 and 2**). The STARLING ensembles show that the microprotein is flexible along its immediate C-terminus, where alanine mutations are tolerated, but the arginine-rich middle segment of the microprotein is more rigid due to the arginines’ repulsive forces. Together these data show that the arginine-rich region is a highly exposed positively charged surface that is critical for its stability and thus likely participates in binding that is necessary for NISM’s activity.

Because mutagenesis impaired the stability of NISM, we next used computational approaches to assess potential direct interactions between NISM and DHX9. DHX9 is a 1,270 amino acid protein that contains multiple folded domains separated by large, disordered regions^54^. DHX9’s folded domains allow for binding to nucleic acids, helicase activity for unwinding primarily R-loops and G quadruplexes, and a transactivation domain for interacting with RNA polymerase II^55,56^. Given the established functions of the folded domains, we hypothesized that the disordered regions of DHX9 could be potential binding sites for the microprotein. To this end, we first used AIUPred to identify four highly disordered regions within DHX9 between known folded domains (**Figure S5C**). Next, we used FINCHES-online’s Intermaps function, which utilizes chemical physics extracted from coarse-grained molecular force field models to predict interactions between disordered regions^57–59^. FINCHES predicted high attraction scores between NISM and three of DHX9’s IDRs, all of which are highly acidic, while the basic C-terminal disordered region was predicted to mostly exhibit repulsive forces (**Figure S5D**). To further support their interaction, we also computed the free energy of DHX9 by itself and in complexation with NISM. In this approach the two protein chains are covalently linked in all combinations, N- to C-terminal, N- to N-terminal, C- to N-terminal, and C to C-terminal. Free energies for these constructs are then calculated accounting for chain connectivity, sequence dependent charge, and non-charge interactions, and the construct with the minimum free energy is chosen. Our earlier work has shown that this approach can reasonably describe experimentally measured relative complexation propensities between two intrinsically disordered proteins^60^. We found that DHX9 complexed with NISM has a lower free energy (−1.5 *kT*) than DHX9 in isolation (0.7 *kT*), thus supporting the interaction between NISM and DHX9. These models, in combination with the IP results, bolster NISM acting as a direct regulator of DHX9 activities.

### NISM overexpression inhibits rRNA synthesis, reduces nucleolar R-loops, and decreases translation

Immunofluorescence experiments and proliferation assays showed that overexpression of NISM-ALFA induces nucleolar shrinkage and downstream stress response activation similarly to ActD (**Figure 2**). Additionally, previous studies have found that many DExD/H-box RNA helicase family members mediate rRNA synthesis and processing^61–63^. Therefore, we measured the effects of both short and long NISM-ALFA overexpression on rRNA synthesis by immunofluorescence. U2OS cells were pulsed with 5-ethynyluridine (5-EU), a ribonucleoside analog which contains an alkyne and therefore allows for conjugation of a fluorophore by click chemistry and quantification of RNA synthesis. Overexpression of NISM-ALFA resulted in decreased nucleolar 5-EU signal compared to control cells, as expected, reflecting less rRNA synthesis (**Figure 5A and 5B**). ActD almost completely blocked rRNA synthesis as expected given its potent inhibition of RNA polymerase I. To further corroborate this effect, 47S pre-rRNA expression was also measured by qRT-PCR. Consistent with the 5-EU incorporation assay, we observed that both NISM-ALFA isoforms significantly inhibited 47S pre-rRNA expression, with ∼40% reduction for short NISM-ALFA, ∼20% reduction for the long isoform, and complete inhibition by ActD treatment (**Figure 5C**). We next wanted to confirm that decreased rRNA synthesis caused by NISM-ALFA overexpression leads to lower translation levels. U2OS cells were pulsed with puromycin and its incorporation in proteins was measured by immunoblot to quantify overall translation levels. Overexpression of both NISM-ALFA isoforms caused a reduction in puromycin incorporation, reflecting inhibition of translation globally (**Figures 5D and 5E**). These results demonstrated that nucleolar stress is activated due to impaired rRNA synthesis and downstream reduction in translation in NISM-ALFA overexpressing cells.

**Figure 5.**
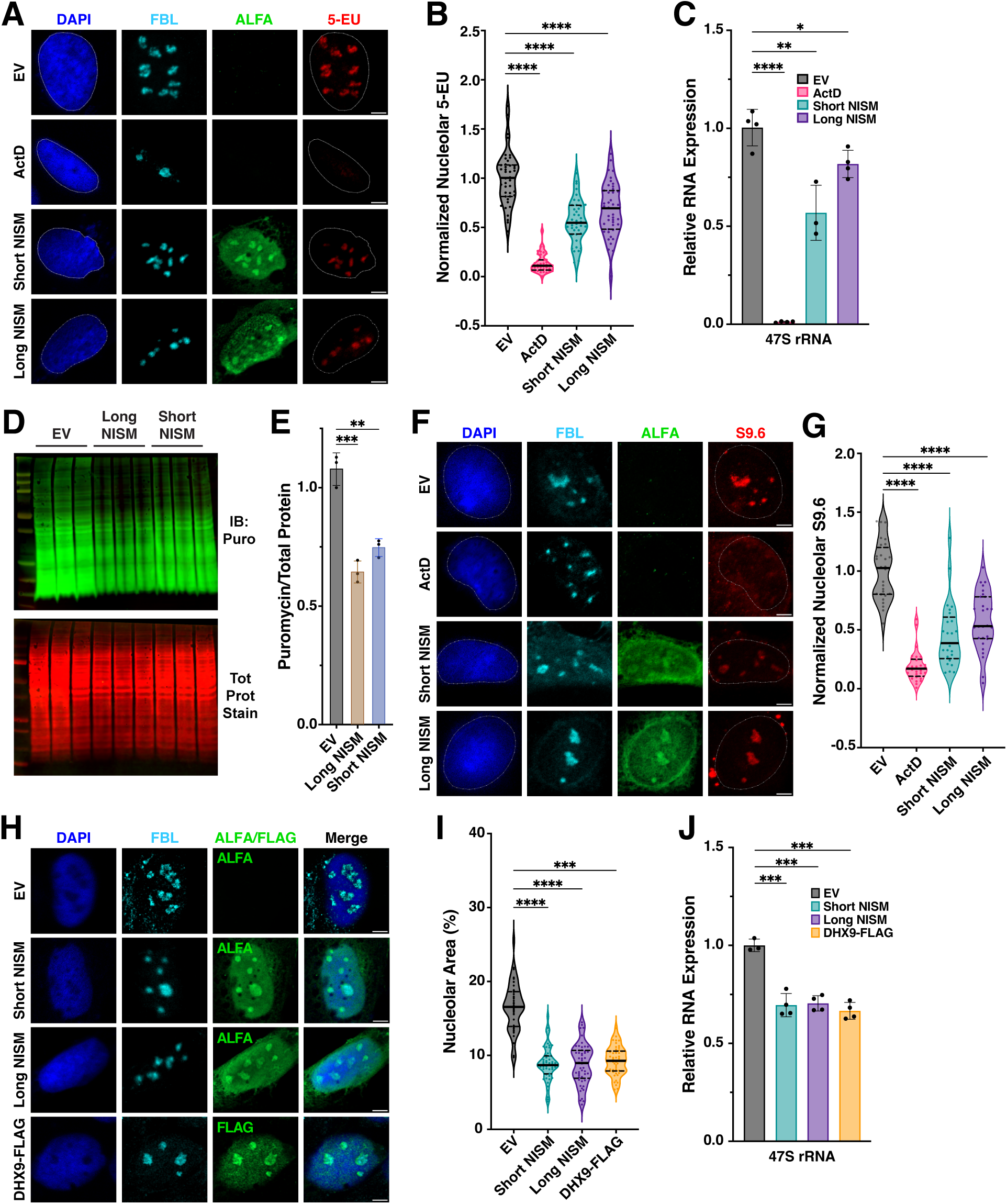
NISM-ALFA overexpression inhibits rRNA synthesis and reduces nucleolar R-loops and overall translation levels. **(A)** Representative immunofluorescence images of FBL (cyan), NISM-ALFA (green), and 5-EU (red) in wild-type (WT) U2OS cells treated with 10 nM ActD or DMSO for 3 h and transfected with empty vector (EV), short NISM-ALFA, or long NISM-ALFA for 48 h. Nuclei were counter-stained with DAPI (blue). Scale bar, 5 µm. **(B)** Quantification of 5-EU staining for cells in (A) represented by violin plots. Statistical comparisons were calculated by Mann Whitney test; n = 40 cells per condition. **(C)** Expression analysis of 47S pre-rRNA by qRT-PCR in cells treated as in (A) represented by bar plots displaying the mean ± SD. Statistical comparisons were calculated by two-tailed Student’s t test; n = 4. **(D)** Immunoblot analysis of puromycin incorporation and total protein loading in WT cells transfected with EV, short NISM-ALFA, or long NISM-ALFA for 48 h. **(E)** Quantification of puromycin staining relative to total protein represented by bar plots displaying the mean ± SD; n = 3. **(F)** Representative immunofluorescence images of FBL (cyan), NISM-ALFA (green), and S9.6 (red) in cells treated as in (A). **(G)** Quantification of S9.6 staining for cells in (D) represented by violin plots. Statistical comparisons were calculated by Mann Whitney test; n = 25 cells per condition. **(H)** Representative immunofluorescence images of FBL (cyan) and NISM-ALFA/DHX9-FLAG (green) in WT cells transfected with EV, short NISM-ALFA, long NISM-ALFA or DHX9-FLAG for 48 h. **(I)** Quantification of nucleolar area for cells in (H) as a percentage of total nuclear area represented by violin plots. Statistical comparisons were calculated by Mann Whitney test; n = 50 cells per condition. **(J)** Expression analysis of 47S pre-rRNA by qRT-PCR in cells treated as in (H) represented by bar plots displaying the mean ± SD. Statistical comparisons were calculated by two-tailed Student’s t test; n = 4. Asterisks indicate: *p < 0.05, **p < 0.01, ***p < 0.001, ****p < 0.0001, ns = not significant. All data are representative of at least two biological replicates.

We next turned our attention to how NISM’s interaction with DHX9 might be implicated in rRNA synthesis inhibition. Eukaryotes have hundreds to thousands of rDNA gene copies located across multiple chromosomes in tandem repetitive clusters, including ∼200–700 in human cells^64,65^. Multiple RNAs beyond rRNA are also transcribed from these gene clusters, including non-coding RNAs (ncRNAs) derived from rDNA intergenic spacer (IGS) regions that are transcribed by both RNA polymerases I and II^66^. Many of these ncRNAs can form R-loops that serve important roles in regulating rRNA synthesis. For instance, R-loops derived from IGS ncRNAs transcribed by RNA polymerase II can serve as “shields” to prevent aberrant expression of sense intergenic non-coding RNAs (sincRNAs) by RNA polymerase I that can disrupt rRNA synthesis^67^. We therefore hypothesized that DHX9’s ability to resolve R-loops and interact with RNA polymerase II may help regulate rRNA synthesis through mediation of ncRNA R-loops. To explore this hypothesis, we assessed the impact of NISM-ALFA overexpression on R-loop formation in nucleoli by immunofluorescence. Overexpression of both NISM-ALFA isoforms resulted in significantly reduced R-loop staining within nucleoli by the S9.6 antibody (**Figure 5F and 5G**). As with 47S pre-rRNA expression, ActD treatment caused a greater reduction in R-loop staining compared to NISM-ALFA overexpression as expected due to inhibition of R-loop formation during transcription^68^. Confirming the specificity of the S9.6 antibody, staining was completely abrogated in cells incubated with RNase H prior to incubation with the antibody (**Figure S6A**). These results suggested that NISM-ALFA overexpression might promote DHX9’s ability to resolve nucleolar R-loops. Indeed, we found that DHX9-FLAG overexpression in U2OS cells also caused a significant decrease in nucleolar area (**Figure 5H-I and S6B-C**) as well as reduced 47S pre-rRNA expression by qRT-PCR (**Figure 5J**).

Importantly, neither NISM-ALFA overexpression nor NISM KO were found to significantly alter DHX9 expression (**Figure S6D and S6E**). Thus, both NISM-ALFA and DHX9 overexpression inhibit rDNA transcription and elicit the nucleolar stress response, further supporting their interaction and functional relationship.

### NISM knockout does not impact rRNA synthesis and translation

Given NISM-ALFA overexpression’s effects on 47S pre-rRNA synthesis and translation, we next examined the impact of NISM KO on these processes. While NISM overexpression inhibited 47S pre-rRNA expression, both NISM KO cell lines showed slight increases in 47S pre-rRNA relative to control cells by qRT-PCR (**Figure S7A**). We also assessed if there were any impacts on pre-rRNA processing. We found modest but significantly increased levels of downstream 18S and 28S rRNAs and no change in 5.8S rRNA levels. To test for any potential downstream effects on ribosome biogenesis, we also measured overall translation by puromycin incorporation. Consistent with there being no reductions in rRNA levels, no change in puromycin labeling was observed in NISM KO cells (**Figures S7B and S7C**). These results showed that while NISM KO induces a p53-dependent cellular stress response and disrupts nucleolar formation (**Figure 3**), overall translation levels did not appear impacted as was observed in NISM-ALFA overexpressing cells.

### NISM is Essential for Proper Nucleolar Formation and Localization of DHX9

After establishing NISM’s ability to interact with DHX9 and their shared regulation of pre-rRNA synthesis, we next examined the impact of NISM knockout on DHX9. We observed that loss of NISM resulted in diffusion of DHX9 throughout the nucleoplasm, even beyond the boundaries of NPM1 staining (**Figures 6A and 6B**). In line with NISM-ALFA’s ability to rescue the cell cycle perturbations observed in the knockout (**Figures 3G and 3H**), we found that NISM-ALFA expression rescued nucleolar formation in both shape and number compared to WT cells, and restored DHX9 nucleolar localization. These results demonstrated that NISM is necessary for nucleolar formation.

**Figure 6.**
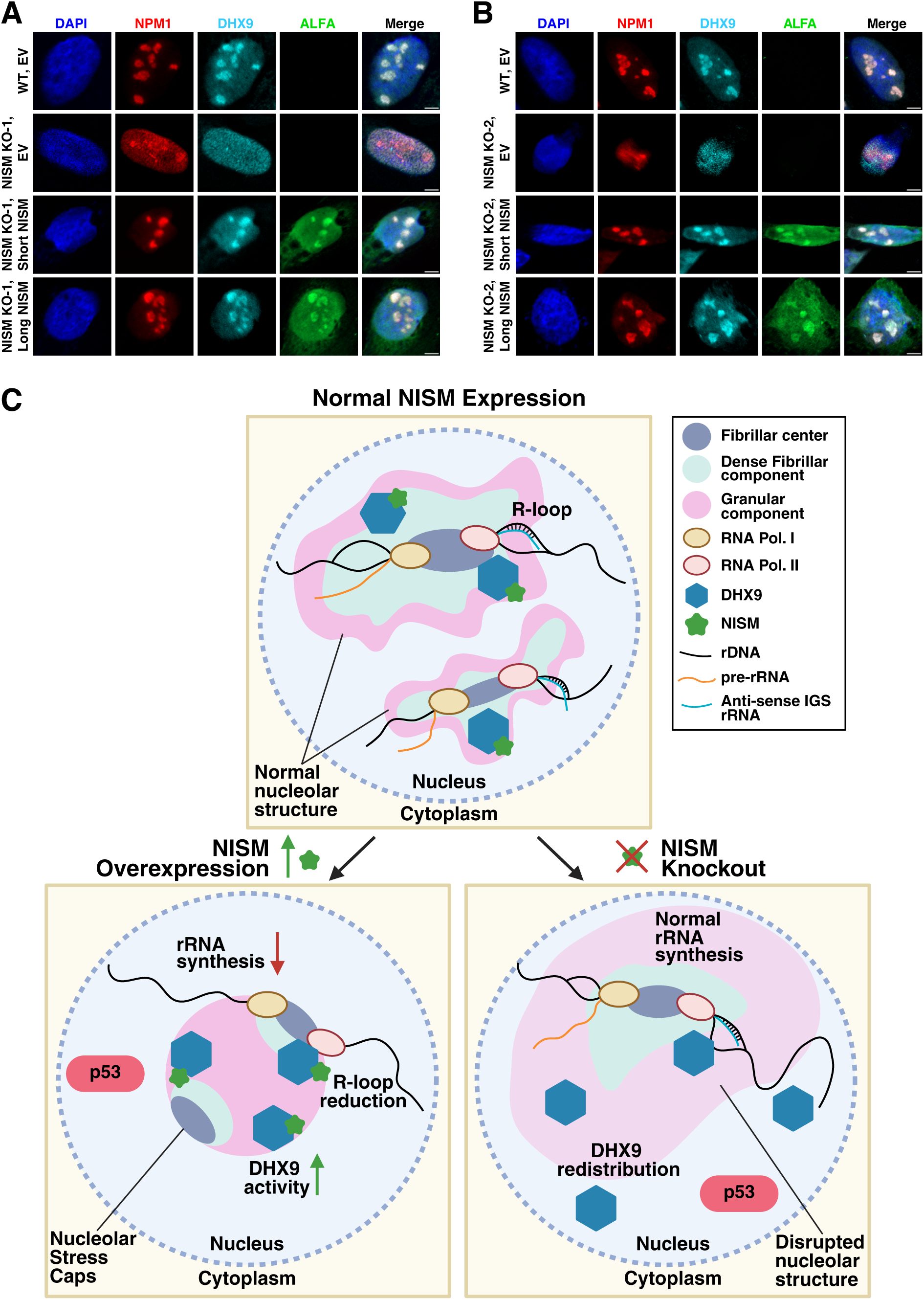
NISM-ALFA rescues nucleolar formation and DHX9 localization in knockout cells. **(A-B)** Representative immunofluorescence images of NPM1 (red), DHX9 (cyan), and NISM-ALFA (green) in wild-type (WT) and NISM knockout (NISM KO) U2OS cells transfected with empty vector, short NISM-ALFA, or long NISM-ALFA for 48 h. Nuclei were counter-stained with DAPI (blue). Scale bar, 5 µm. All data are representative of at least two biological replicates. **(C)** Proposed model for how NISM perturbation regulates nucleolar dynamics. In WT U2OS cells, NISM interacts with DHX9 to enhance its LLPS propensity, thus allowing for proper formation of nucleoli. Synthesis and processing of rRNA, nucleolar R-loop homeostasis, and all steps of ribosome biogenesis proceed normally. When NISM is overexpressed, DHX9 nucleolar phase separation is increased, in turn causing a reduction in nucleolar R-loops, impaired rRNA synthesis, and the formation of stress caps at the outer edges of shrunken, rounded nucleoli. These effects lead to p53 stabilization, cell cycle checkpoint activation, and suppressed cell proliferation. In contrast to overexpression, NISM knockout causes disruption of the nucleolar structure, thus resulting in dispersion of nucleolar proteins and redistribution of DHX9 to the nucleoplasm. While rRNA synthesis and overall translation levels are not significantly impacted, p53 is stabilized and cell proliferation is strongly suppressed.

Nucleolar assembly has been shown to result from liquid-liquid phase separation (LLPS) of interacting RNA and protein components^69–71^. Thus, we next assessed NISM’s potential to promote LLPS using the computational tools ParSe v2^72^ and catGRANULE 2.0^73^. ParSe analyzes protein sequences for disordered regions and whether these regions can promote phase separation. ParSe predicted a disordered region for the C-terminal 24 residues of both short and long NISM, similar to AIUPred (**Figure S2B**), but did not predict this region to promote phase separation (**Supplemental Data 3**). Similarly, catGRANULE 2.0, which uses physiochemical properties and AlphaFold to predict LLPS potential at single amino acid resolution, reported relatively low LLPS scores for NISM (**Figure S8A-C, Supplemental Data 3**). However, while NISM is unlikely to promote LLPS of the nucleolus, DHX9 showed strong potential. ParSe reported multiple disordered regions that can likely undergo LLPS for DHX9, particularly its long C-terminal disordered region. catGRANULE also reported a high LLPS propensity score for DHX9’s C-terminus. Notably, the prediction that NISM can directly interact with each of DHX9’s disordered domains except its C-terminus (**Figure S5D**) still allows for DHX9 to drive LLPS via its C-terminal disordered region. These results suggested that NISM’s ability to regulate nucleolar formation likely depends on regulation of DHX9’s LLPS propensity through their interaction.

Given that both NISM and DHX9 are highly enriched in charged residues, we next used charge patterning analysis to determine how complexation with NISM impacts DHX9’s ability to interact with itself and thus its propensity LLPS. For this purpose, we constructed a Sequence Charge Decoration Matrix (SCDM) of DHX9 in the presence and absence of NISM. SCDM is a collection of patterning metrics that arise from the sequence of charged amino acids in an intrinsically disordered protein. Specifically, the elements of SCDM (i,j) quantify the electrostatic interaction contribution to the ensemble average distance between two residues^74,75^. This comparison showed that DHX9 in the presence of NISM has more intra-chain attraction than in the absence of NISM (**Figure S8D**). Previous studies have shown that attractive intra-chain interactions can be correlated to inter-chain attraction and hence higher propensity for LLPS^76,77^. Thus, these analyses indicate that NISM’s interaction with DHX9 can promote phase separation.

Collectively, our data support a model in which NISM impacts nucleolar dynamics by binding to DHX9 and promoting its phase separation (**Figure 6C**). Without NISM present, DHX9 is unable to promote proper nucleolar formation by LLPS and it localizes throughout the nucleus. This triggers p53 activation and suppressed proliferation. When NISM is overexpressed, DHX9 localizes to nucleoli, but its phase separation may be enhanced such that its activity is altered. This leads to R-loop reduction and inhibited 47S pre-rRNA synthesis, which also leads to p53 activation and suppressed proliferation.

## DISCUSSION

The discovery and characterization of NISM adds to our growing understanding of how microproteins can regulate the function of larger proteins and protein complexes to impact cellular biology^1,2,78^. Here, we show that NISM is a novel regulator of nucleolar formation and rRNA synthesis. Based on our empirical data, computational analyses, and mathematical modeling, we posit that NISM’s essentiality for proper nucleolar formation and its impact on 47S pre-rRNA synthesis are both dependent on NISM’s ability to bind to the DExH-box RNA helicase DHX9 and promote its LLPS activity. Supporting this model, DHX9 was among the highest detected nucleolar prey proteins in the IP-MS datasets (**Supplemental Data 2**) and co-localizes with DHX9 within nucleoli (**Figures 4E and 6A-B**). In addition, we found that DHX9’s C-terminal disordered region is predicted to have a high LLPS propensity, which is promoted by complexation with NISM, while NISM itself is not predicted to promote LLPS (**Figure S8** and **Supplemental Data 3**). Finally, NISM can potentially interact directly with each of DHX9’s disordered regions except the C-terminus and therefore would not block its ability to interact with other biomolecules to promote LLPS (**Figure S5**). These data together point to NISM acting as a modulator of DHX9 to enable its nucleolar activities.

The ability of DExD/H-box family helicases to regulate phase separation as well as their involvement in nucleolar biology are well known^79–83^. For instance, DDX18 is essential for nucleolar phase separation and nucleolar substructure organization through interactions with NPM1 as well as RNAs^84^. Similarly, DDX10 has been shown to be necessary for nucleolar formation via two of its IDRs, and it has a role in 18S rRNA maturation in mouse embryonic stem cells^62^. RNA helicases can also impact nucleolar organization by mediating interactions between the nucleolus and heterochromatin, such as the *Drosophila* DEAD-box helicase Pitchoune^85^. DHX9, too, been shown to have roles nucleolar formation and rRNA synthesis. In vascular smooth muscle cells, for example, DHX9 knockdown was found to disrupt nucleolar structure^19^. Interestingly, this same study also reported that the lncRNA *MIAT* interacts with DHX9 and is necessary for its localization to the nucleolus, similar to what we observe for NISM. Moreover, *MIAT* promotes DHX9’s interaction with PARP1, which was also enriched in NISM-ALFA IP-MS experiments (**Supplemental Data 1**). However, knockdown of *MIAT* caused a reduction in 47S pre-rRNA, differentiating its activity from NISM’s. These examples all demonstrate the importance of DExD/H-box helicases in LLPS driven formation of the nucleolus and their subsequent functions in rRNA synthesis and processing, thus supporting how NISM likely regulates these processes through its interaction with DHX9.

In considering NISM overexpression-induced nucleolar stress, we hypothesize that NISM’s role in promoting DHX9 LLPS is also involved in these processes. First, NISM overexpression caused a modest reduction in both 47S pre-rRNA levels (**Figures 5A-C**) and nucleolar R-loops (**Figures 5F and 5G**). Nucleolar R-loops are key regulators of rDNA transcription in the nucleolus, though they have contrasting effects depending on which R-loops are made.

Previous studies investigating the loss of *SETX*^86^ and *GPATCH4*^87^, have found that accumulation of nucleolar R-loops impairs pre-rRNA synthesis. However, specific RNA polymerase II transcribed R-loop shields within rDNA IGS regions are critical for pre-rRNA synthesis, and loss of these shields disrupt nucleolar structure^67^. Our results would suggest that these R-loop shields, or similar rDNA transcription promoting R loops, are inadvertently resolved by DHX9 due to enhancement of its activity by NISM overexpression. Supporting this possibility, DHX9 overexpression mimicked the effects of NISM overexpression on nucleolar shrinkage and rDNA transcription (**Figures 5H-J**). While DHX9 does not have a reported role in nucleolar R-loop resolution, it has been shown that the R-loop interactome is enriched for proteins with long IDRs and high propensity for LLPS, and thus may facilitate the formation of membrane-less R loop foci^88^. Indeed, LLPS has been suggested to play a role in the resolution of double-strand break-induced R-loops by DHX9^89^. In addition, DHX9 has previously been found to bind to and promote the processing of IGS-rRNA into pRNA to establish nucleolar heterochromatin during embryonic stem cell differentiation^49^. DHX9’s ability to interact with IGS-rRNA might hint at its ability to also interact with IGS R-loops. It is also possible nucleolar R-loops are reduced due to impaired formation, as opposed to their resolution. In this scenario, enhanced DHX9 activity could lead to broad transcriptional inhibition due to increased heterochromatin formation or physical blocks of RNA polymerases that in turn blocks the synthesis of pre-rRNA and IGS ncRNAs, both of which contribute to R-loop formation. Investigating the mechanism of NISM overexpression on nucleolar R-loop resolution and formation, and DHX9’s involvement, further would be of interest for future studies.

Curiously, we observed that 47S pre-rRNA synthesis and overall translation levels were not significantly impacted by the loss of nucleolar structure resulting from NISM KO (**Figure S7**). These results suggested that while p53 is activated and cell proliferation is suppressed (**Figure 3**), this is not a direct consequence of any negative impacts on translation levels. However, we cannot rule out that while overall translation levels are unchanged in the knockout cells, aspects of ribosome biogenesis are compromised such that there are additional free subunits or ribosomal proteins. For instance, knockdown of the nucleolar protein PAK1IP1 caused an increase in soluble RPL5 and RPL11, which in turn caused p53 stabilization and accumulation of cells in the G0/G1 phase, similar to what we observed in NISM KO cells^90^. It is also worth noting that DHX9 depletion has been shown to cause G0/G1 accumulation in mouse fibroblasts, which is thought to be due to loss of DHX9 localization with CIZ1 in the nucleolus^52^.

Finally, beyond the characterization of NISM, this study also strengthens support for the additional thousands of putative arginine-rich microproteins having roles in RNA metabolism (**Figure S1B**). NISM’s role in regulating nucleolar formation joins it with NoBody as a microprotein regulator of MLO formation and RNA metabolism^8,91^. It now also joins MINAS-60^92^ and alt-LAMA3^93^ as regulators of ribosome biogenesis. However, NISM is unique compared to these other microproteins due to its significantly smaller size, 36 amino acids for NISM versus 68 for NoBody, 130 for MINAS-60, and 148 for alt-LAMA3, as well as the proposed mechanism for how it impacts MLOs and ribosome biogenesis. For example, NoBody has been shown to directly promote LLPS to regulate P-body formation^91^, while NISM represents, to our knowledge, the first example of a microprotein that can regulate the LLPS propensity of another protein.

Additionally, NISM’s role in ribosome biogenesis differs from that of MINAS-60, which regulates pre-60S assembly, and alt-LAMA3, which regulates 47S pre-rRNA synthesis through interaction with PeBoW. We expect that continued investigation of arginine-rich microproteins will deepen our understanding of ribosome biogenesis and other RNA-related processes.

## METHODS

### Materials

| REAGENT or RESOURCE | SOURCE | IDENTIFIER |
| --- | --- | --- |
| **Antibodies** |  |  |
| FluoTag®-X2 anti-ALFA LI-COR IRDye 800CW | NanoTag Biotechnologies | Cat# N1502-Li800-L; RRID: AB_3075985 |
| Anti-β-Actin antibody, Mouse Monoclonal Clone AC-15 | Sigma-Aldrich | Cat# A1978 |
| Recombinant sdAb anti-ALFA Guinea Pig Fc-fusion | NanoTag Biotechnologies | Cat# N1584; RRID: AB_3075999 |
| NPM1 Monoclonal Antibody (FC-61991) | Thermo Fisher Scientific | Cat# 32-5200; RRID: AB_2533084 |
| UBF-1 Polyclonal Antibody | Thermo Fisher Scientific | Cat# PA5-36153; RRID:<br>AB_2553396 |
| RPA194 Antibody (C-1) | Santa Cruz Biotechnology | Cat# sc-48385; RRID:<br>AB_675814 |
| Fibrillarin (C13C3) Rabbit Monoclonal Antibody | Cell Signaling Technology | Cat# 2639; RRID:<br>AB_2278087 |
| p53 (DO-7) Mouse Monoclonal Antibody | Cell Signaling Technology | Cat# 48818; RRID:<br>AB_2713958 |
| DHX9 (E6M4Q) Rabbit Monoclonal Antibody | Cell Signaling Technology | Cat# 71286 |
| beta-Tubulin (9F3) Rabbit Monoclonal Antibody | Cell Signaling Technology | Cat# 2128; RRID:<br>AB_823664 |
| Histone H3 Antibody | Cell Signaling Technology | Cat# 9715; RRID:<br>AB_331563 |
| DDX5 (D15E10) Rabbit Monoclonal Antibody | Cell Signaling Technology | Cat# 9877; RRID:<br>AB_2616008 |
| Normal Guinea Pig Control IgG | SinoBiological | Cat# CR4 |
| ANTI-FLAG® M2 antibody, Mouse monoclonal | Sigma-Aldrich | Cat# F3165; RRID:<br>AB_259529 |
| Anti-DNA-RNA Hybrid Antibody, clone S9.6 | Sigma-Aldrich | Cat# MABE1095; RRID:<br>AB_2861387 |
| HA-Tag (C29F4) Rabbit Monoclonal Antibody (BSA and Azide Free) | Cell Signaling Technology | Cat# 86124; RRID:<br>AB_1549585 |
| Normal Rabbit IgG | Cell Signaling Technology | Cat# 2729; RRID:<br>AB_1031062 |
| HDAC5 (D1J7V) Rabbit Monoclonal Antibody | Cell Signaling Technology | Cat# 20458; RRID:<br>AB_2713973 |
| Anti-Puromycin Antibody, clone 12D10 | Sigma-Aldrich | Cat# MABE343; RRID:<br>AB_2566826 |
| IRDye® 800CW Goat anti-Mouse IgG Secondary Antibody | LI-COR Biosciences | Cat# 926-32210; RRID:<br>AB_621842 |
| IRDye® 800CW Goat Anti-Rabbit IgG | LI-COR Biosciences | Cat# 926-32211; RRID:<br>AB_621843 |
| IRDye® 680RD Goat anti-Mouse IgG Secondary Antibody | LI-COR Biosciences | Cat# 926-68070; RRID:<br>AB_10956588 |
| Goat anti-Rabbit IgG (H+L) Highly Cross-Adsorbed | Thermo Fisher Scientific | Cat# A-11073; RRID:<br>AB_2534117 |
| Secondary Antibody, Alexa Fluor™ 488 |  |  |
| Goat anti-mouse IgG, FSD™ 594 | BioActs | Cat# RSA1195 |
| Goat anti-mouse IgG, FSD™ 594 | BioActs | Cat# RSA1275 |
| **Biological Samples** |  |  |
| Short NISM-ALFA | GenScript | N/A |
| Long NISM-ALFA | GenScript | N/A |
| pSpCas9(BB)-2A-Puro (PX459) V2.0 | Addgene | Ran et al.<sup>94</sup>; Addgene Plasmid #62988 |
| Homo sapiens DExH-box helicase 9 (DHX9), transcript variant 1, mRNA. pcDNA3.1-C-(k)DYK | GenScript | Cat# OHu107233 |
| Short NISM Alanine Mutants, pcDNA3.1+ | GenScript | N/A |
| **Chemicals, peptides, and recombinant proteins** |  |  |
| KpnI-HF | New England Biolabs | Cat# R3142 |
| XbaI | New England Biolabs | Cat# R0145 |
| Corning® 500 mL DMEM (Dulbecco's Modified Eagle's Medium) | Corning | Cat# 10-013-CV |
| GenClone® Fetal Bovine Serum | GenClone | Cat# 25-550 |
| McCoy's 5A Medium | ATCC | Cat# 30-2007 |
| Lipofectamine™ 2000 Transfection Reagent | Thermo Fisher Scientific | Cat# 11668027 |
| Lipofectamine™ LTX Reagent with PLUS™ Reagent | Thermo Fisher Scientific | Cat# 15338100 |
| Opti-MEM™ I Reduced Serum Medium | Thermo Fisher Scientific | Cat# 31985070 |
| 1M Tris-HCl, pH 7.4. 1000 mL, Sterile | Teknova | Cat# T1074 |
| Sodium Chloride, 5 Molar, Ionic Strength Adjustor for Potassium ISE | RICCA Chemical Company | Cat# 7227-16 |
| EDTA 0.5M Solution pH 8.0 | Research Products International | Cat# E14000 |
| Triton™ X-100 Stock Solution (20% v/v) | Cayman Chemical | Cat# 600217 |
| cOmplete ULTRA EDTA-free Protease Inhibitor Cocktail | Roche | Cat# 05892791001 |
| 4X Bolt™ LDS Sample Buffer | Thermo Fisher Scientific | Cat# B0007 |
| 2-Mercaptoethanol | Sigma-Aldrich | Cat# 63689 |
| 20X Bolt™ MES SDS Running Buffer | Thermo Fisher Scientific | Cat# B0002 |
| Power Blotter Pre-cut Membranes and Filters, PVDF, regular size | Thermo Fisher Scientific | Cat# PB9320 |
| Glycine, Free Base, 100 Grams | Research Products International | Cat# G36050 |
| Tricine | Sigma-Aldrich | Cat# T0377 |
| Methanol, ACS Grade | StatLab | Cat# 40380-1 |
| Intercept® (TBS) Blocking Buffer | LI-COR Biosciences | Cat# 927-60001 |
| Poly-D-lysine | Thermo Fisher Scientific | Cat# A3890401 |
| DMSO, sterile filtered | Tocris Biosciences | Cat# 3176 |
| Actinomycin D | Sigma-Aldrich | Cat# A1410 |
| PBS, pH 7.4 | Thermo Fisher Scientific | Cat# 10010023 |
| 16% Formaldehyde, Methanol-Free | Cell Signaling Technology | Cat# 12606 |
| Apex TBS, 20X | Apex Biosearch | Cat# 18-236B |
| Tween 20 [Polyoxyethylenesorbitan Monolaurate] | Research Products International | Cat# P20370 |
| Normal Goat Serum | Jackson Immuno Research | Cat# 005-000-121 |
| ProLong™ Gold Antifade Mountant with DNA Stain DAPI | Thermo Fisher Scientific | Cat# P36935 |
| Corning® 100 mL Trypsin EDTA 1X | Corning | Cat# 25053CI |
| Ethanol | Gold Shield | Cat# 64-17-5 |
| RNase A, DNase and protease-free (10 mg/mL) | Thermo Fisher Scientific | Cat# EN0531 |
| Propidium Iodide | Sigma-Aldrich | Cat# P4170 |
| Crystal violet solution | Sigma-Aldrich | Cat# V5265 |
| 1M HEPES | Thermo Fisher Scientific | Cat# 15630-080 |
| Potassium Chloride (White Crystals) | Thermo Fisher Scientific | Cat# BP366 |
| Magnesium Chloride Hexahydrate | Thermo Fisher Scientific | Cat# BP214 |
| Pierce™ IP Lysis Buffer | Thermo Fisher Scientific | Cat# 87787 |
| Sepharose™ Protein A/G | Rockland Immunochemicals | Cat# PAG50-00-0002 |
| Trichloroacetic acid | Sigma-Aldrich | Cat# T6399 |
| Acetone, ACS Grade | Thermo Fisher Scientific | Cat# S25120 |
| Urea, 1kg | Apex Biosearch | Cat# 20-205 |
| TCEP-HCl | Apex Biosearch | Cat# 18-217 |
| 2-chloroacetamide | Sigma-Aldrich | Cat# 22790 |
| 1M TEAB | Thermo Fisher Scientific | Cat# 90114 |
| Promega Sequencing Grade Modified Trypsin, Lyophilized | Promega Corporation | Cat# PRV5111 |
| Formic Acid, 88% | Thermo Fisher Scientific | Cat# S25331 |
| Empore 47 mm SDB-XC extraction disks | CDS Analytical | Cat# 2240 |
| Acetonitrile | Sigma-Aldrich | Cat# 34851 |
| Trifluoroacetic acid, 99+%, for HPLC | Thermo Fisher Scientific | Cat# AC432291000 |
| TRI Reagent | Zymo Research Corporation | Cat# R2050-1-200 |
| Chloroform – isoamyl alcohol mixture | Supelco | Cat# 25666 |
| Glycogen, molecular biology grade | Thermo Fisher Scientific | Cat# FERR0561 |
| Isopropanol, 99.5%, for molecular biology, DNase, RNase and Protease free | Thermo Fisher Scientific | Cat# 327270010 |
| Molecular Biology Grade Water | Corning | Cat# 46-000-CM |
| RNase H | New England Biolabs | Cat# M0297L |
| Gold Biotechnology Inc Bovine Serum Albumin (BSA), Fraction V, Protease Free | Gold Biotechnology Incorporated | Cat# A-420 |
| Puromycin Dihydrochloride | Research Products International | Cat# P330200 |
| Revert™ 700 Total Protein Stain Kit | LI-COR Biosciences | Cat# 92611016 |
| BbsI | New England Biolabs | Cat# R0539 |
| T4 DNA ligase | New England Biolabs | Cat# M0202 |
| NEB® 5-alpha Competent E. coli (High Efficiency) | New England Biolabs | Cat# C2987 |
| Puromycin dihydrochloride from Streptomyces alboniger | Sigma-Aldrich | Cat# P8833 |
| QuickExtract DNA extraction solution | Lucigen | Cat# QE09050 |
| Agarose | Apex Biosearch | Cat# 20-102GP |
| Apex TBE Buffer 10X | Apex Biosearch | Cat# 20-131 |
| SYBR™ Gold Nucleic Acid Gel Stain (10,000X Concentrate in DMSO) | Thermo Fisher Scientific | Cat# S11494 |
| **Critical commercial kits** |  |  |
| Q5 Site-Directed Mutagenesis kit |  |  |
| BCA Protein Assay Kit with BSA Protein Standard | Prometheus Protein Biology Products | Cat# 18-440 |
| WST-1 Cell Proliferation Assay Kit | Cayman Chemical | Cat# 10008883-5 |
| RevertAid First Strand cDNA Synthesis Kit | Thermo Fisher Scientific | Cat# K1621 |
| KiCqStart SYBR Green qPCR ReadyMix | Sigma-Aldrich | Cat# KCQS01 |
| Click-iT™ RNA Alexa Fluor™ 594 Imaging Kit | Thermo Fisher Scientific | Cat# C110330 |
| 2X Phusion™ Plus PCR Master Mix | Thermo Fisher Scientific | Cat# F631S |
| GeneJET PCR Purification kit | Thermo Fisher Scientific | Cat# K0701 |
| ZR Plasmid Miniprep-Classic Kits | Zymo Research Corporation | Cat# D4054 |
| **Software** |  |  |
| FlowJo (version 10.10.0) | BD Biosciences |  |
| Fiji | Schindelin et al.<sup>95</sup> |  |
| Cell Profiler | Carpenter et al.<sup>96</sup> |  |
| Zen 3.12 | Carl Zeiss AG3 |  |
| Fragpipe V19 | Yu et al.<sup>97</sup> |  |
| Prism 10.3.1 | GraphPad |  |

### Experimental model

#### Cell Culture

Cell lines used in this study were purchased from ATCC. HEK293T (CRL-3216) cells were cultured in DMEM with 4.5 g/L glucose, L-glutamine & sodium pyruvate (Corning, 10-013-CV) supplemented with 10% (vol/vol) FBS (GenClone, 25-550). U2OS (HTB-96) cells were cultured in McCoy’s 5A medium with 1.5 mM L-glutamine and 2200 mg/L sodium bicarbonate (ATCC, 30-2007) supplemented with 10% (vol/vol) FBS. Cells were incubated at 37°C with 5% CO_2_ in a humidified environment. Cells were regularly tested for mycoplasma contamination.

### Method details

#### Expression constructs and cell transfection

Short and Long NISM with C-terminal ALFA and DHX9 with C-terminal FLAG were synthesized and cloned into pcDNA3.1(+) (Thermo Fisher Scientific, V7920) by GenScript. Short and Long NISM with C-terminal HA was generated in-house using the Q5 Site-Directed Mutagenesis kit (New England Biolabs, E0554S) per manufacturer protocol to substitute the DNA sequence coding for ALFA to HA. All primers for Q5 cloning were designed using the NEBaseChanger™ tool.

The C-terminal Short NISM alanine mutant constructs (Ala 2-7, Ala 8-13, Ala 14-18, Ala 19-23, Ala 24-36) was synthesized by GenScript and cloned into pcDNA3.1(+) between the KpnI and XbaI restriction sites. Alanine mutants 2-12 and 13-23 were synthesized as DNA fragments and cloned into pcDNA3.1(+) in house using double-digest restriction enzyme cloning with restriction enzymes KpnI-HF (New England Biolabs, R3142) and 20U XbaI (New England Biolabs, R0145).

For overexpression experiments, constructs were transfected into HEK293T and U2OS cells using Lipofectamine 2000 (Thermo Fisher Scientific, 11668027) and Lipofectamine LTX with PLUS reagent (Thermo Fisher Scientific, 15338100), respectively, with 2 μg plasmid DNA and 6 μL transfection reagent per sample (1:3 DNA:lipofecamine) in 6-well plates (GenClone, 25-105). Transfection complexes were prepared in 400 μL Opti-MEM (Gibco, 31985070) and incubated for 20 min before adding directly to cells at a confluency of 50% for HEK293T and 80% for U2OS cultured in 2 mL media (1:5 transfection mix:media). For immunofluorescence (IF) experiments, fresh media was replaced 24 h after transfection. Cells were analyzed 48 h after transfection for all experiments except for cell cycle and p53 experiments, where they were analyzed 72 h after transfection.

#### Immunoblot analysis

Cells were lysed using ice cold lysis buffer containing 50 mM Tris (Teknova, T1074), 150 mM NaCl (RICCA Chemical Company, 7227-16), 1 mM EDTA (Research Products International, E14000) and 1% Triton X-100 (Cayman Chemical, 600217) supplemented with 1x cOmplete protease inhibitor cocktail (Roche, 05892791001). Crude lysate was clarified by centrifugation at 13,000 x g for 10 min at 4°C and protein concentration was measured using a BCA assay (Prometheus, 18-440) for SDS-PAGE.

Samples for SDS-PAGE and immunoblotting were prepared with SDS protein loading dye (Thermo Fisher Scientific, B0007) to a final concentration of 1X and 2.5% BME (Sigma-Aldrich, 63689) and boiled at 95°C for 5 min. 25 μg protein were resolved on a 4-12% Bis-Tris gel (Thermo Fisher Scientific,, NW04120BOX) using MES-SDS running buffer (Thermo Fisher Scientific, B0002) at 180 V for 32 min using the PowerEase Touch 120W power supply (Thermo Fisher Scientific, PS0120). Proteins were transferred to PVDF membranes (Thermo Fisher Scientific, PB9320) with the Power Blotter semi-dry transfer system (Thermo Fisher Scientific, PB0012) at 1.3 A for 10 min using transfer buffer containing 0.34 M Tris, 0.26 M Glycine (Research Products International, G36050), 0.14 M Tricine (Sigma-Aldrich, T0377), and 2% (vol/vol) methanol (StatLab, 40380-1). Membranes were blocked in blocking buffer (LI-COR Biosciences, 927-60001) at room temperature for 1 h.

The following primary antibody dilutions were used for each experiment described in the sections below. ALFA immunoblots were incubated with 800CW conjugated anti-ALFA nanobody (NanoTag Biotechnologies, N1502-Li800-L) diluted 1:500 in blocking buffer. β-Actin immunoblots were incubated with mouse anti-B Actin primary antibody (Sigma-Aldrich, A1978) diluted 1:2500 in blocking buffer. Histone H3 and β-tubulin immunoblots were incubated with rabbit monoclonal anti-H3 (Cell Signaling Technology, 9715) and rabbit polyclonal anti-β-tubulin (Cell Signaling Technology, 2128) antibodies diluted 1:1000 in blocking buffer. DHX9 and DDX5 immunoblots were incubated with rabbit monoclonal anti-DHX9 (Cell Signaling, 71286) and anti-DDX5 (Cell Signaling, 9877) diluted 1:500 in blocking buffer. FLAG immunoblots were incubated with mouse monoclonal anti-FLAG (Sigma-Aldrich, F3165) Membranes were washed with TBS (Apex Bioresearch, 18-236B) supplemented with Tween (Research Products International, P20370) 0.1% 3 times at room temperature for 5 min each wash. Primary antibody was detected by 800CW conjugated anti-mouse (LI-COR Biosciences, 926-32210 or anti-rabbit (LI-COR Biosciences, 926-32211) IgG secondary antibody diluted 1:10,000 in blocking buffer at room temperature for 1 h and washed 3 times with TBST 0.1% before imaging using the Odyssey CLx imager (LI-COR Biosciences, 9140). For immunoblots of DHX9-FLAG, 680RD conjugated anti-mouse (LI-COR Biosciences, 926-32211) was used to detect FLAG.

#### Immunofluorescence imaging

4-well chamber slides (NEST Scientific USA, 230104) were treated with Poly-D-lysine (Gibco, A3890401) according to the manufacturer protocol prior to seeding 20,000 HEK293T or 30,000 U2OS cells in 400 μL of media to each well of the chamber slide. The following day, cells were transfected with 0.4 μg plasmid DNA per condition and after 24 h, fresh media was replaced per well. 48 h after transfection, or 72 h for p53 experiments, cells were treated with either DMSO (Tocris, 3176) or Actinomycin D (Sigma-Aldrich, A1410) at 10 nM for 3 h. Cells were washed once with PBS (Gibco, 10010-023) and fixed with 4% methanol-free PFA (Cell Signaling, 12606P) diluted in PBS at room temperature for 10 min. Cells were washed with PBS 3 times and permeabilized with 0.2% Triton X-100 (Cayman Chemical, 600217) diluted in PBS at room temperature for 10 min with gentle rocking. They were then washed three times in PBST 0.1% and blocked at room temperature for 1 h with blocking buffer containing 3% (vol/vol) goat serum (Jackson Immuno Research, 005-000-121) diluted in PBST 0.1%. Primary antibodies were incubated at 4°C overnight with recombinant heavy-chain anti-ALFA guinea pig antibody (NanoTag Biotechnologies, N1584) diluted 1:500, HA rabbit monoclonal antibody (Cell Signaling Technology, 86124) diluted 1:500, FLAG mouse monoclonal antibody diluted 1:1000 (Sigma-Aldrich, F3165), NPM1 mouse monoclonal antibody (Thermo Fisher Scientific, 32-5200) diluted 1:500, UBF-1 rabbit polyclonal antibody (Thermo Fisher Scientific, PA5-36153) diluted 1:200, RPA194 mouse monoclonal antibody (Santa Cruz Biotechnology, sc-48385) diluted 1:200 and FBL rabbit monoclonal antibody (Cell Signaling Technology, 2639) diluted 1:400 in blocking buffer for nucleolar morphology experiments. For p53 experiments, p53 mouse monoclonal antibody (Cell Signaling Technology, 48818) was diluted 1:500 in blocking buffer.

For DHX9 localization experiments, DHX9 rabbit monoclonal antibody (Cell Signaling, 71286) was diluted 1:500 in blocking buffer. After primary antibody incubation, cells were washed 3 times with PBST 0.1% and incubated with secondary antibodies at room temperature for 1 h, protected from light. Recombinant guinea pig anti-ALFA was detected using goat anti-guinea pig IgG AF488 (Thermo Fisher Scientific, A-11073). Mouse monoclonal antibodies were detected using goat anti-mouse IgG FSD594 (BioActs, RSA1195) and rabbit monoclonal antibodies were detected using goat anti-rabbit IgG FSD680 (BioActs, RSA1275). Cells were washed 3 times with PBST 0.1% and dried before mounting using medium containing DAPI (Thermo Fisher Scientific, P36935) and sealing with a glass coverslip (Fisher, 12-541-019). Samples were imaged using a Zeiss LSM900 Airyscan confocal microscope (Carl Zeiss AG) with a 63×1.4NA oil immersion DIC M27 objective. Chromatic aberrations were corrected in the Zen software using translation channel alignment with the KNN algorithm and images were analyzed using FIJI software.

#### Fluorescence image quantification

To quantify nucleolar area, NPM1 channel was converted to an 8-bit grayscale image in FIJI and the nucleoli were thresholded. Individual cells were then segmented using the DAPI channel and the % area of thresholded NPM1 signal was measured to quantify nucleolar area per segmented cell. For DHX9-FLAG immunofluorescence quantification, the nucleolar area was quantified using FBL.

To quantify p53 signal intensity, 20X stitched images of each sample were captured and signal intensity was measured using CellProfiler. To segment individual U2OS cells by their nuclei, the identify primary objects module was used, with object diameter set between 150-to-400-pixel units. Global thresholding was performed using the Otsu method and clumped objects were separated by object shape. To smoothen nuclei edge segmented in the identify primary objects module, the identify secondary objects module was applied, with global thresholding performed using the Otsu method. Pixel expansion was set to 100 for the secondary segmentation. The signal intensity of the p53 or ALFA channel for segmented nuclei was then measured in 16-bit and normalized by maximum signal intensity per image and plotted on a scale between 0 and 1 as arbitrary units. In NISM overexpression experiments, microprotein positive cells were thresholded using the ALFA channel to quantify p53.

To quantify the NPM1 coefficient of variation (*CV*), individual cells were segmented using the DAPI channel. The mean NPM1 intensity (*μ* _(NPM1)_) and standard deviation SD (σ) was measured per cell. *CV* was calculated by dividing the SD (σ) by the mean.

To quantify nucleolar S9.6 intensity, the S9.6 channel was converted to 16-bit in FIJI. Nucleoli were then thresholded using the FBL channel and segmented with the region of interest (ROI) tool. This ROI was then used to measure mean intensity of S9.6 in the nucleolus.

#### WST-1 assay

U2OS cells were seeded at a density of 300,000 cells/well in a 6 well plate. The following day, 2 μg plasmid DNA were transfected for each sample. 24 h after transfection, cells were washed with PBS and seeded into 96-well plates (GenClone, 25-109) in quadruplicate at a density of 3,000 cells/well in 100 μL of media. For untransfected cells, they were first seeded into a 6 well plate and then passaged into into 96-well plates after 24 h. After an additional 24 h, cells were treated with either DMSO or 0.5 nM Actinomycin D. Cell proliferation was measured using a WST-1 assay kit (Cayman Chemical, 10008883-5) according to the manufacturer’s protocol and absorbance at 450 nm at 37°C was measured for days 0-3 using a Victor Nivo multimode microplate reader (Revvity, HH35000500). For each day, wells containing media only were also treated with WST-1 reagents and measured as a blank background signal. Statistics was performed on technical replicates on day 3 of proliferation.

#### Cell cycle analysis by flow cytometry

For cell cycle assessment of U2OS cells, DNA content was measured by flow cytometry. 300,000 U2OS cells were seeded per well into each well of a 6-well TC plate. 24 h after seeding, 2 μg plasmid DNA were transfected in duplicates for each sample. The following day, transfected cells were trypsinized (Corning, 25053CI) and seeded into 10-cm TC dishes (GenClone, 25-202). 72 h after transfection, cells were treated with DMSO or Actinomycin D at 10 nM for 3 h. For cell cycle assessment HDAC5-MP KO cells, 750,000 cells/well were seeded into 6-cm TC dishes (GenClone, 25-260) and transfected in duplicates with 4 μg plasmid DNA for rescue condition with 72 h overexpression. After drug treatment, cells were washed with PBS and 1 million cells were counted for cell cycle analysis. Cells were fixed with 1 mL ice-cold 70% ethanol (Gold Shield, 64-17-5) in a dropwise manner to a final volume of 1 mL, and the cells were resuspended by gentle pipetting. After fixation, cells were centrifuged at 130 x g and washed three times with ice-cold PBS. After the 3^rd^ wash, the supernatant was carefully removed, and cells were resuspending in 100 μL PBS containing RNase A (Thermo Fisher Scientific, EN0531) at a concentration of 100 μg/mL at room temperature for 5 min. After RNase A treatment, DNA was stained by adding 400 μL PBS containing propidium iodide (Sigma Aldrich, P4170) at a concentration of 50 μg/mL at room temperature for 30 min, protected from light. Cells were analyzed by flow cytometry on a BD LSRFortessa X-20 (Becton-Dickson) using the following voltage acquisition settings on a linear scale; FSC: 340V, SSC: 193V and PI: 338V. 10,000 cells per samples were acquired at a slow flow rate. Data was analyzed using FlowJo v10.10.0 and fitted to the Watson (Pragmatic) model.

#### Colony formation assay

For the colony formation assay WT or NISM KO U2OS cells were seeded into Poly-D-lysine treated 6-well plates. Prior to cell count, the cell suspension was passed through a 40 µm cell strainer (Corning, 431750) to ensure a single cell suspension. For each cell line, cells were seeded in technical triplicates at a density of 1,500 cells/well. Cells were grown for two weeks with media changes every 3 days. To stain colonies, cells were washed once with ice-cold PBS and fixed with ice-cold methanol for 15 min at 4°C. Cells were then incubated with 1% stock crystal violet (Sigma-Aldrich, V5265) diluted in 20% methanol for 30 min at room temperature. Cells were then washed 3 times with water.

To quantify colonies, FIJI was used. Images of each well were converted to 8-bit and the thresholding tool was used to detect stained colonies. The colonies were then quantified by % area within each well.

#### Nuclear fractionation

U2OS cells were grown to 80-90% confluency in 15-cm TC dishes (GenClone, 25-203) using 20 mL media volume and transfected with 20 μg plasmid DNA in triplicates per condition. 48 h after transfecting, cells were washed with ice-cold PBS and gently scraped into 500 μL nuclear isolation buffer containing 10 mM HEPES (Gibco, 15630-080), 100 mM KCl (Fisher BioReagents, BP366), 5 mM MgCl_2_ (Fisher BioReagents, BP214) and 0.05% Triton X-100 supplemented with 1X cOmplete protease inhibitor cocktail. Cells were left to lyse on ice for 20 min with periodic vortexing. Cells were centrifuged at 4°C at 700 x g for 5 min and the supernatant containing the cytosolic fraction was collected for quality assessment. The remaining nuclear pellet was washed once with 500 μL of nuclear isolation buffer and centrifuged at 4°C at 700 x g. The supernatant was removed, and the nuclear pellet was lysed with 600 μL Pierce IP lysis buffer (Thermo Fisher Scientific, 87787) on ice for 20 min with periodic vortexing. After lysis, the crude lysate was clarified by centrifugation at 17k x g for 20 min at 4°C and protein concentration was measured using a BCA assay. A fraction of the nuclear lysate was collected for SDS-PAGE and the remaining nuclear lysate was used for IP, normalizing input to 500 μg at a concentration of 1 mg/mL. Purity of the nuclear fraction was then assessed by SDS-PAGE and immunoblot using B-tubulin rabbit monoclonal antibody (Cell Signaling, 2128) and Histone H3 rabbit polyclonal antibody (Cell Signaling, 9715).

#### Nuclear lysate co-immunoprecipitation

First, 20 μL of Sepharose protein A/G resin (Rockland Immunochemicals, PAG50-00-0002) was washed with 1 mL cold TBST 0.1%. Next, nuclear lysates were pre-cleared by incubation with 0.5 μg control guinea pig IgG (SinoBiological, CR4) and the pre-washed resin under rotation at 4°C for 1 h. After pre-clearance, lysates were incubated with 0.5 μg recombinant guinea pig anti-ALFA antibody for IP at 4°C with rotation overnight. The following day, 20 μL pre-washed Sepharose protein A/G resin was used to pulldown recombinant guinea pig anti-ALFA IgG at 4°C with rotation for 2 h. After incubation, the samples were spun down at 1,000 x g for 2 min at 4°C and the unbound fraction was collected for SDS-PAGE. Sepharose resin was washed with 1 mL ice-cold TBST 0.1% three times under rotation at 4°C for 10 min per wash. . For proteomics, captured proteins were eluted from resin using 50 μL of elution buffer containing 0.2 M glycine, pH 2.6 at room temperature with shaking at 900 rpm for 10 min in a thermomixer (Eppendorf, 05-412-503). Samples were centrifuged at 1,000 x g and supernatant containing elution fraction 1 was collected and neutralized with 50 μL Tris pH 9.0 neutralization buffer. A second elution fraction was collected, neutralized, and combined with the first elution fraction for a total volume of 200 μL. 30 μL of this eluate was collected for SDS-PAGE and the remaining 170 μL volume was processed further for mass spectrometry.

#### Mass spectrometry sample preparation

The eluate for mass spectrometry analysis was supplemented with 30 μL of trichloroacetic acid, TCA, (Sigma Aldrich, T6399) to a final concentration of 15% (vol/vol) TCA and stored at - 20°C overnight to precipitate proteins. To pellet proteins, samples were centrifuged at 17k x g for 30 min at 4°C. The supernatant was partially removed, leaving ∼20 μL volume, and the protein pellet was washed twice with 1 mL ice-cold acetone (Fisher, S25120), with centrifugation at 17k x g for 20 min at 4°C after each wash. The supernatant was removed, and the protein pellet was dried for 5 min before redissolving in 12.5 μL 8M urea (Apex Bioresearch, 20-205).

Proteins were reduced by adding 1.5 μL 100 mM tris(2-carboxyethyl) phosphine hydrochloride, TCEP, (Apex Bioresearch, 18-217) to a final concentration of 5 mM and incubated at 37°C for 30 min with shaking at 900 rpm in a thermomixer. Proteins were then alkylated by adding 1.4 μL 500 mM 2-chloroacetamide (Sigma-Aldrich, 22790) to a final concentration of 50 mM and incubated at 37°C for 30 min, shaking at 900 rpm, protected from light. 30.5 μL 100 mM TEAB (Thermo Fisher Scientific, 90114) and 1 μL trypsin (Promega Corporation, PRV5111) at a concentration of 0.5 μg/μL was added to digest proteins at 37°C overnight with shaking at 900 rpm. The following day, formic acid (Thermo Fisher Scientific, S25331) was added to a final concentration of 10% (vol/vol) to quench the digest.

Digested samples were desalted using Empore 47 mm SDB-XC extraction disks (CDS Analytical, 2240) packed into 200 μL pipette tips. The disks were pre-equilibrated with wetting buffer containing 80:20:0.1% water: acetonitrile (Sigma-Aldrich, 34851): trifluoroacetic acid (Thermo Fisher Scientific, AC432291000) followed by wash buffer containing 98:2:0.1% water: acetonitrile: trifluoroacetic acid. Digest samples were then loaded on the extraction disks and washed twice with wash buffer before eluting with elution buffer containing 60:40:0.1% water: acetonitrile: trifluoroacetic acid. All volumes used for equilibrating, washing, or eluting were 30 μL with centrifugation at 700 x g for 2 min between each step. Eluted peptides were dried in a speed vac and redissolved in acetonitrile and formic acid. Digested and cleaned peptides were analyzed by LC MS/MS using an UltiMate 3000 RSLC (Thermo Fisher, San Jose, CA) coupled to an Orbitrap Fusion Lumos mass spectrometer (Thermo Fisher, San Jose, CA). Peptides were separated by reverse-phase on a 50 cm × 75 μm I.D. Acclaim® PepMap RSLC column using gradients of 4% to 22% acetonitrile at a flow rate of 300 nL/min (solvent A: 100% water, 0.1% formic acid; solvent B: 100% acetonitrile, 0.1% formic acid) for 67 min (total acquisition time: 90 min). For each MS/MS acquisition, duty cycles consisted of one full Fourier transform scan mass spectrum (375–1800 m/z, resolution of 120,000 at m/z 400) followed by data-dependent MS2 acquired at top speed in the linear ion trap for 3 s at 100% normalized AGC. Ions detected in MS1 with 2+ or greater charge were selected and subjected to HCD fragmentation (NCE 30%) in MS2 and resulting ions were detected in the linear ion trap in ‘Rapid’ mode with normalized AGC target 200%. Ions that were selected for MS2 were dynamically excluded from selection for 30 sec.

#### Mass spectrometry analysis

Raw files were analyzed using FragPipe v19 and searched against a custom database combining the single protein per gene UniProt human database (UP000005640, April 2022) and Ribo-seq detected microproteins (**Supplemental Data 1**) using LFQ search parameters with the following settings: Precursor mass tolerance was set to ± 50 ppm and the fragment mass tolerance was set to 600 ppm. Semi-enzymatic trypsin cleavage after lysine or arginine except before proline was used with a maximum of 2 missed cleavages. Peptide length was set to range 6-50 and mass range was set to 600-6000 Da. Methionine oxidation and N-terminal acetylation were set as variable modifications with maximum occurrences of 3 and 1, respectively, and carbamidomethylation of cysteine was set as a fixed modification. False discovery rates (FDR) were employed using the target-decoy approach with 1% FDR set for proteins. MS1 quantification was performed using IonQuant^98^ with match between runs and MaxLFQ options selected, and intensities were normalized across runs. For downstream analysis, each set of triplicate samples were grouped according to their experimental conditions and protein interactions were scored using SAINTexpress by comparing spectral counts of the bait condition to an EV control. Hits were filtered using a SAINT score ≥ 0.8 and an average spectral count between 3 replicates ≥ 5. Common prey proteins between short and long NISM-ALFA were then analyzed by STRING^99^ to identify interactions among the prey proteins using the associated webserver tool (https://string-db.org/). Network edges were set to confidence, with the line thickness indicating the strength of data support between protein interactions. Interaction networks were then clustered using k-means clustering with the number of clusters set to 3.

For validation of IP, 35 μg of input nuclear lysate samples and 30 μL of IP samples were analyzed by SDS-PAGE and immunoblot. Elution was performed by boiling beads in 60 μL 2X SDS with 2.5% BME. DHX9 was detected using 1:500 anti-DHX9 antibody diluted in LiCOR blocking buffer, incubated at 4°C overnight with gentle shaking. IP of DHX9 was performed using 0.5 μg DHX9 antibody with the same conditions described previously, pre-clearing with 0.5 μg control Rabbit IgG (Cell Signaling, 2729).

#### qRT-PCR

For qRT-PCR, U2OS cells were seeded at a density of 300,000 cells/well in 6-well plates and transfected in duplicate using 2 μg of plasmid DNA for 48 h overexpression. Total RNA was extracted by lysing cells with 1 mL of TRI Reagent (Zymo Research Corporation, R2050-1-200) per well. Lysates were incubated at room temperature for 5 min before adding 200 μL chloroform (Supelco, 25666), and samples were vigorously vortexed and incubated at room temperature for 3 min. Samples were centrifuged at 12,000 x g for 15 min and the aqueous phase containing RNA was collected. 1 μL of glycogen (Thermo Fisher Scientific, FERR0561) and 500 μL of isopropanol (Thermo Fisher Scientific, 327270010) were added to the aqueous phase and vigorously vortexed. Samples were then stored at -20°C overnight to precipitate RNA. The following day, samples were centrifuged at 4°C at 12k x g for 15 min to pellet RNA. The supernatant was removed and the RNA pellet was washed with 1 mL ice-cold nuclease free 70% ethanol and centrifuged 4°C at 12,000 x g for 15 min. This wash was repeated once, and the RNA pellet was dried for 5 min before resuspending in 25 μL of nuclease free water (Corning, 46-000-CM). RNA concentration was measured using a Nanodrop (Thermo Scientific, ND-2000C). 1 μg of total RNA was used for cDNA synthesis using the RevertAid First Strand cDNA Synthesis Kit (Thermo Scientific, K1621) per manufacturer protocol with random hexamer primers. cDNA was then diluted 1:30 and 20 μL qPCR reaction volumes were prepared in technical duplicates per sample using the KiCqStart SYBR Green qPCR ReadyMix (Sigma Aldrich, KCQS01). qPCR was performed using the QuantStudio 7 Flex standard 96-well format (Thermo Scientific, 4485688) with the following cycle settings: 95°C for 3 min, followed by 40 cycles of 95°C for 10 s to 60°C for 1 min, and a final melt curve stage at 95°C for 15 s to 60°C for 1 min and to 95°C for 15 s. The following primers sets were used at a final concentration of 450 nM and were synthesized by Integrated DNA Technologies: 47S rRNA—F: 5′-GAACGGTGGTGTGTCGTTC-3′, R: 5′-CGTCTCGTCTCGTCTCACTC-3′; 18S rRNA—F: 5′-AAACGGCTACCACATCCAAG-3′, R: 5′-CCTCCAATGGATCCTCGTTA-3′; 28S rRNA—F: 5′-CCTCCAATGGATCCTCGTTA-3′, R: 5′-ATGACGAGGCATTTGGCTAC-3′; 5.8S rRNA—5′-CTCTTAGCGGTGGATCACTC-3′, R: 5′-GACGCTCAGACAGGCGTAG-3′; GAPDH—F: 5′-GAAGAGAGAGACCCTCACTGCTG-3′, R: 5′-ACTGTGAGGAGGGGAGATTCAGT-3′. Data were analyzed using the 2(–ΔΔCt) method with GAPDH as a reference gene for normalization.

#### 5-EU nascent RNA labeling assay

For click chemistry labeling of cellular RNA synthesis, the Click-iT™ RNA Alexa Fluor™ 594 Imaging Kit (Thermo Fisher Scientific, C10330) was used. Untransfected or transfected cells were grown to 70% confluency on a Poly-D-lysine treated 4-well chamber slide in 400 μL media and treated with 10 nM Actinomycin D or DMSO for 3 h. After drug treatment, 400 μL of warmed media containing 2 mM 5-EU was added to each well to a final concentration of 1 mM. After 1 h incubation with 5-EU, the cells were washed with PBS and fixed at room temperature for 10 min with methanol-free 4% PFA. Fixed cells were washed once with PBS and permeabilized with 0.2% Triton X-100 diluted in PBS at room temperature for 15 min. After permeabilization, the click reaction was performed per manufacturer protocol, using 400 μL of reaction volume per well, and all steps were performed protected from light. Following the click reaction step, immunofluorescence was performed starting with the blocking step, described previously to co-stain FBL or ALFA.

To analyze mean nucleolar 5-EU (*μ* _(nucleolar EU)_), the mean nucleoplasm EU (*μ* _(nucleoplasm EU)_) intensity per cell was measured in FIJI and used for background subtraction from total mean nuclear EU (*μ* _(total nuclear EU)_) (Eq. 1). The EU channel was then converted to 16-bit grayscale and nucleolar EU was quantified by thresholding. The nucleus was segmented individually per cell using the DAPI channel and this segmentation was applied to the EU channel. Thresholded EU signal within individually segmented nuclei were quantified by measuring the mean gray value. All measurements were then normalized to the mean of the negative control.

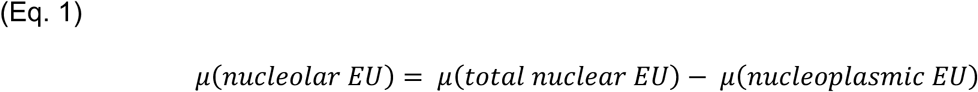

#### DNA-RNA Hybrid Immunofluorescence Imaging Assay

Transfected U2OS cells seeded into 4-well chamber slides and overexpressing HDAC5-MP for 48 h were treated with 10 nM Actinomycin D or DMSO for 3 h. After drug treatment, cells were washed once with PBS. After thoroughly aspirating the PBS from each well, cells were quickly fixed with 200 μL of ice-cold methanol at -20°C for 5 min. After fixing, cells were rehydrated with PBS and incubated for 5 min per wash for a total of 3 washes. Cells were quenched with 200 μL 0.1 M glycine prepared fresh at room temperature for 5 min, followed by 3 washes with PBS for 5 min each wash. Next, cells were treated with 0.5% Triton X-100 diluted in PBS for 7 min. As a control, RNase H was used to digest DNA-RNA hybrids for each condition using 400 μL RNase H (New England Biolabs, M0297L) diluted 1:50 in 1X RNase H buffer at 37°C for 5 h. Cells were washed twice with PBS and blocked with blocking buffer containing 2% BSA (GoldBio, A-420) and 1% FBS diluted in PBS at room temperature for 1 h.

Cells were incubated with primary antibody directed against DNA-RNA Hybrids using the S9.6 mouse monoclonal antibody (Sigma-Aldrich, MABE1095) diluted 1:250 in 3% goat serum, along with ALFA and FBL antibodies diluted as described previously above. The same secondary antibodies were used per primary antibody previously described.

#### Puromycin incorporation assay

For puromycin incorporation experiments, 750,000 U2OS cells were seeded in 6-cm cell culture plates for both overexpression and NISM KO conditions. The following day, U2OS cells were transfected with 4 μg of plasmid DNA for 48 h prior to labeling. For knockout cells, cells were labeled the following day after seeding into 6-cm dishes. Cells were treated with puromycin (Research Products International, P330200) at a working concentration of 4 μg/mL in complete media for 30 min. Cells were then lysed with 200 μL of lysis buffer described previously and protein concentration was measured using a BCA assay.

For each sample, 40 μg of protein was resolved by SDS-PAGE. Proteins were then transferred to a PVDF membrane overnight at 5V with Towbin buffer made up of 25 mM Tris, 192 mM glycine, and 20% methanol. After transfer, membranes were stained for total protein using the Revert™ 700 Total Protein Stain (LI-COR Biosciences, 92611016) per manufacturer protocol. Total protein was imaged using the Odyssey CLx imager. Membranes were then washed three times with TBST 0.1% for 10 min per wash and blocked with LI-COR blocking buffer for an hour. To detect puromycin labeled proteins, membranes were incubated overnight at 4°C with mouse monoclonal anti-puromycin antibody (Sigma-Aldrich, MABE343) diluted 1:1000 in blocking buffer. After primary antibody incubation, membranes were washed three times with TBST 0.1% and incubated with secondary antibody described previously.

#### Generation of clonal NISM knockout cells

HDAC5-MP KO U2OS cell lines were generated using an all-in-one CRISPR/Cas9 plasmid system delivered to cells via transient transfection. Single gRNAs targeting exon 1 of the HDAC5 gene were designed using the Custom Alt-R CRISPR-Cas9 gRNA tool and synthesized by Integrated DNA Technologies. DNA oligos for the gRNA sequences (Sense: 5′-CACCGCAAAGATGGAGGAGCCGTCG-3′; Antisense: 5′-AAACCGACGGCTCCTCCATCTTTGC-3′) were annealed and cloned into the pSpCas9(BB)-2A-Puro (PX459) V2.0 (Addgene, 62988) using the BbsI restriction enzyme (New England Biolabs, R0539) and T4 DNA ligase (New England Biolabs, M0202). Ligated constructs were transformed into high efficiency 5-alpha Competent E. coli (New England Biolabs, C2987) and colonies were selected, amplified and mini-prepped (Zymo, D4054) for Sanger Sequencing.

U2OS cells were transfected with 2 μg of plasmid in 6-well plates and after 24 h, transfected cells were selected using 2.0 μg/mL puromycin (Sigma-Aldrich, P8833) for 2 days. Selected cells were then expanded, and clonal cell lines were sorted into 96-well plates using the BD FACS Aria Fusion flow cytometer (Becton-Dickson). Clonal cell lines were expanded for 3 weeks with media changes every 3-4 days. At ∼80% confluency, a fraction of each isolated clonal cell line was harvested to freeze down at an early passage number, and the remaining cells were used for genotyping.

Genomic DNA was extracted from cells using QuickExtract DNA extraction solution (Lucigen, QE09050) and genomic knockouts were screened by PCR amplification of the genomic locus using the following set of genotyping primers: F: 5′-GGTCTGGGTCTATTTTTAGCTCCGG-3′, R: 5′-GGGTCCGGGGAAGATGGGAT-3′ in aa Analytik-Jena Biometra TRIO 48 thermal cycler (Analytik Jena, 846-2-070-723). 40 μL PCR reaction mixes were prepared using 2X Phusion™ Plus PCR Master Mix (Thermo Scientific, F631S) to amplify 100 ng of template DNA with forward and reverse primers added to a final concentration of 0.5 μM. The following PCR cycle was used: 95°C for 3 min, followed by 35 cycles of 98°C for 30 s to 58°C for 30 s to 72°C for 30 s, and a final extension stage at 72°C for 4 min. WT DNA produced a PCR product of 240 bp and PCR products were resolved on a 4% agarose (Apex Bioresearch, 20-102GP) gel stained with SYBR gold (Thermo Fisher Scientific, S11494) at 105 V for 2 h using TBE buffer (Apex Bioresearch, 20-131). PCR amplicons from clonal KO cell lines that displayed homozygous indels bands were purified using the GeneJET PCR Purification kit (Thermo Scientific, K0701) and sequenced by Sanger sequencing with the forward genotyping primer. Expression of the HDAC5 mRNA was validated in WT or KO cells using qPCR with primers spanning exon-exon junctions designed using Primer-BLAST: Exons 2 & 3: F: 5′-GGCATGAACTCTCCCAACGA-3′, R: 5′-GGCTTCACCTCCACTGTCAC-3′; Exons 8 & 9: F: 5′-TGTTGAGATCACAGGTGCCG-3′, R: 5′-GCTGAGGGAGCATCTCAGTG-3′.

Expression of HDAC5 at the protein level was also validated by SDS-PAGE and immunoblot using the HDAC5 rabbit monoclonal antibody (Cell Signaling, 20458) following IP of HDAC5 from 500 μg of total protein from WT or NISM KO U2OS cells.

#### Computational Analyses of NISM structure, disorder, confirmational ensemble, binding, and LLPS propensity

Short and long NISM structures were predicted using ESMfold^28^, which was run using default settings in the Google Colab environment created by ColabFold^27^ (ESMFold.ipynb). AIUpred^21^ disorder predictions for NISM were run via the associated webserver (https://aiupred.elte.hu/) with the settings “Only disorder” and “Default smoothing” selected. STARLING^13^ was used to predict the confirmational ensembles for short and long NISM. STARLING was run using default settings to generate 200 conformations for each NISM isoform. The starling2xtc command was then used to make trajectory files for each ensemble. Finally, trajectory files were covered to movies to visualize the different conformations using PyMOL. FINCHES^57^ was used to predict interactions between each of DHX9’s IDR and short NISM using the Intermaps tool on the FINCHES webserver (https://finches-online.com/). LLPS propensity for NISM and DHX9 were predicted using ParSe v2^72^ (https://stevewhitten.github.io/Parse_v2_web/) and catGRANNULE 2.0^73^ (https://tools.tartaglialab.com/catgranule2) with default settings via their associated webservers.

#### Computing Free Energy of Complexation

To calculate the binding free energy of two sequences, we covalently linked the Short NISM protein with the DHX9 protein at their ends. For every pair, we consider four possible combinations: i) C-terminus end of sequence A linked to N-terminus end of sequence B, ii) C-terminus end of sequence A linked with C-terminus end of sequence B, iii) N-terminus end of sequence A linked with C-terminus end of sequence B, and iv) N-terminus end of sequence A linked with N-terminus end of sequence B. Next, we used Equation 2 below derived by our group previously to compute the free energy of an IDP with arbitrary sequence^75,100^.

(Eq. 2)

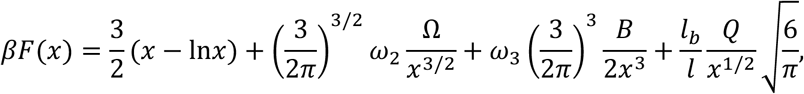

where, Ω, 𝑄, and 𝐵 contain details of the composite sequence, and are given by

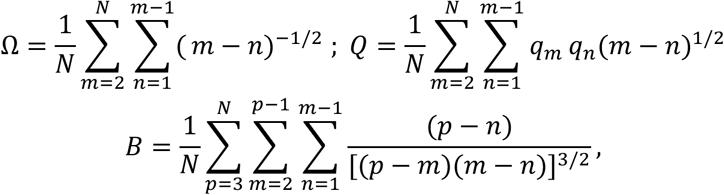

𝑥 is a measure of chain dimension, and the free energy is minimized with respect to 𝑥 to determine the most likely conformation and free energy at this conformation. The electrostatic component of the free energy is determined from the sequence dependent 𝑄 term computed by using 𝑞 = −1 for amino acid residues D, E and 𝑞 = 1 for amino acid residues R, K and zero for everything else. The non-electrostatic interaction parameter 𝜔_2_ is sequence dependent and its value is determined from a physics-based machine learning algorithm trained against coarse-grained simulation^101^ and three body excluded volume parameter 𝜔_3_ = 0.2. For each composite sequence the minimum free energy is computed for these different constructs and the lowest value from these minimums was chosen as the free energy of the complex.

#### Computing Sequence Charge Decoration Matrix

Sequence Charge Decoration Matrix (𝑆𝐶𝐷𝑀*_ij_*) computes electrostatic interactions between any two amino acids 𝑖 and 𝑗, and can be further used to compute ensemble average distance 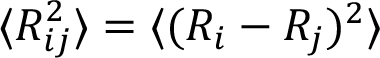 = ⟨(𝑅*_i_*− 𝑅*_j_*)^2^⟩ between 𝑖, 𝑗, where, 𝑅*_i_* and 𝑅*_j_* are position vectors of some representative atoms of residue 𝑖 and 𝑗. Specifically, 𝑆𝐶𝐷𝑀 is defined as Equation 3^74,75^,

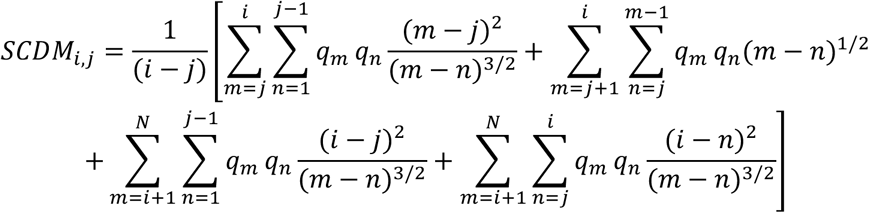

with, 𝑞*_i_* = −1 for amino acids 𝐷, 𝐸, 𝑞*_i_* = 1 for amino acids 𝑅, 𝐾 and zero otherwise. Thus, 𝑆𝐶𝐷𝑀 gives an electrostatic patterning map which quantifies interaction that dictates conformation.

## Statistical Analyses

Statistics were performed using the two-tailed unpaired Student’s t-test or Mann Whitney test, and a p value of < 0.05 was used as a cut off for significance. For all p values, the level of significance was indicated as follows: *, p < 0.05; **, p < 0.01; ***, p < 0.001; ****, p < 0.0001. Values greater than 0.05 were considered not significant and indicated by ns. All error bars are plotted with mean ± SD in GraphPad Prism (Version 10.3.1).

## ACKNOWLEDGEMENTS

This work was supported by the National Institutes of Health (R35GM157126 to T.F.M., R01GM138901 to K.G., and R37CA252081, R21AI185033, and R21ES036190 to R.B.). The content is solely the responsibility of the authors and does not necessarily represent the official views of the National Institutes of Health. R.B. was supported by a Research Scholar Grant (RSG-24-1249960-01-DMC) from the American Cancer Society. Salary support for P.O. was provided by a California Institute for Regenerative Medicine (CIRM) stem cell biology training grant (TG2-01152) and an EMBO Postdoctoral fellowship (ALTF 213-2023). We thank members of the Martinez lab as well as Drs. Alex Holehouse, Kyoko Yokomori, Andrej Luptak, Cholsoon Jang, and Ivan Marazzi for thoughtful discussions and suggestions. We also thank Drs. Clinton Yu and Lan Huang of the UCI High-End Mass Spectrometry Facility and Chao Family

Comprehensive Cancer Center (P30CA062203) for assistance with mass spectrometry proteomics experiments, Dr. Vanessa Scarfone and Ms. Pauline Nguyen of the UCI Flow Cytometry Core for assistance with cell cycle distribution experiments, and Dr. Allia Fawaz of the UCI Microscope Imaging Core for assistance with imaging experiments. NISM model in Figure 6 was created with BioRender.com.

## RESOURCE AVAILABILITY

### Lead Contact

Thomas F. Martinez

### Materials Availability

All materials, including NISM KO cells and expression constructs, are available upon request.

### Data and Code Availability

The mass spectrometry proteomics data have been deposited to the ProteomeXchange Consortium via the PRIDE^102^ partner repository with the dataset identifier PXD074472. No new code was written for this study.

## AUTHOR CONTRIBUTIONS

T.F.M. conceived, organized, and managed project implementation; K.C., D.H., J.H., J.W., N.H., and P.O.M. conducted experiments. T.F.M., R.B., K.C., D.H., G.T., J.H., J.W., N.H., H.E.D., and P.O.M. analyzed the results; L.H. and K.G. performed free energy and SCDM calculations.

T.F.M., K.G., and R.B. supervised the study and secured funding; T.F.M. wrote the original draft of the manuscript and all authors participated in review and editing of the manuscript.

## SUPPLEMENTAL INFORMATION

**Document S1.** Figures S1-S8.

**Supplemental Data 1.** Compilation of all smORFs called translated by RibORF 1.0 and RiboCode in previously generated HEK293T, HeLa-S3, and K562 Ribo-seq datasets^10^ and filtered lists of microproteins containing different ARMs.

**Supplemental Data 2.** SAINTexpress results for significantly enriched prey proteins identified in nuclear IP-MS experiments of short and long NISM-ALFA.

**Supplemental Data 3.** ParSe v2 and catGRANULE 2.0 results for short and long NISM and DHX9.

**Supplemental Videos 1 and 2.** STARLING conformational ensembles predicted for short and long NISM.

