## Supplemental Figures for "*HDAC5*-encoded Microprotein NISM Mediates Nucleolar Formation and Ribosomal RNA Synthesis"

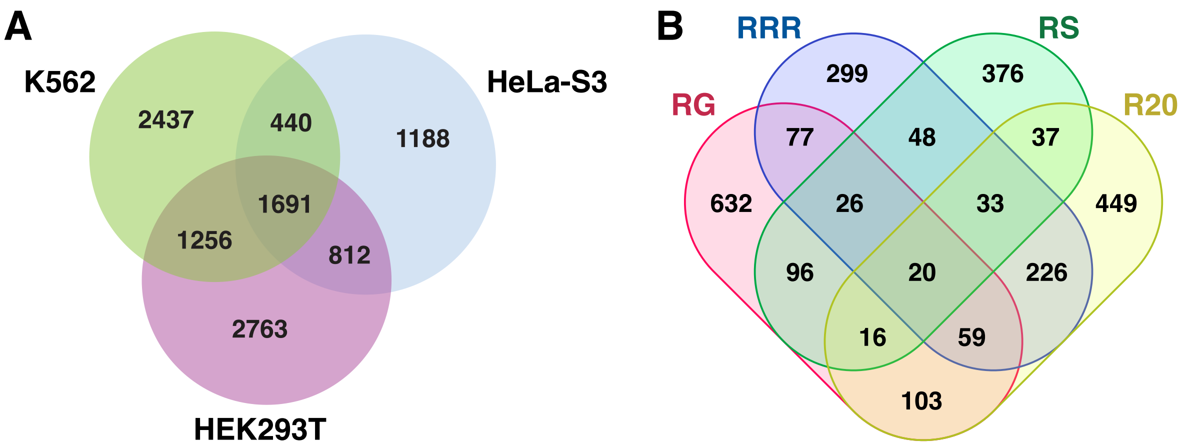


**Figure S1. Thousands of putative microproteins contain arginine-rich motifs, related to Figure 1. (A)** Venn diagram showing the overlap of smORFs called translated by both RibORF v1.0 and RiboCode in previously published Ribo-seq datasets collected from HEK293T, HeLa-S3, and K562 cells[10]. **(B)** Venn diagram showing the overlap of putative microproteins containing different arginine-rich motifs. RG denotes microproteins with two or more RG motifs, RS denotes two or more RS motifs, RRR denotes two or more RRR motifs, and R20 denotes microproteins with arginine representing at least 20% of the amino acid sequence.

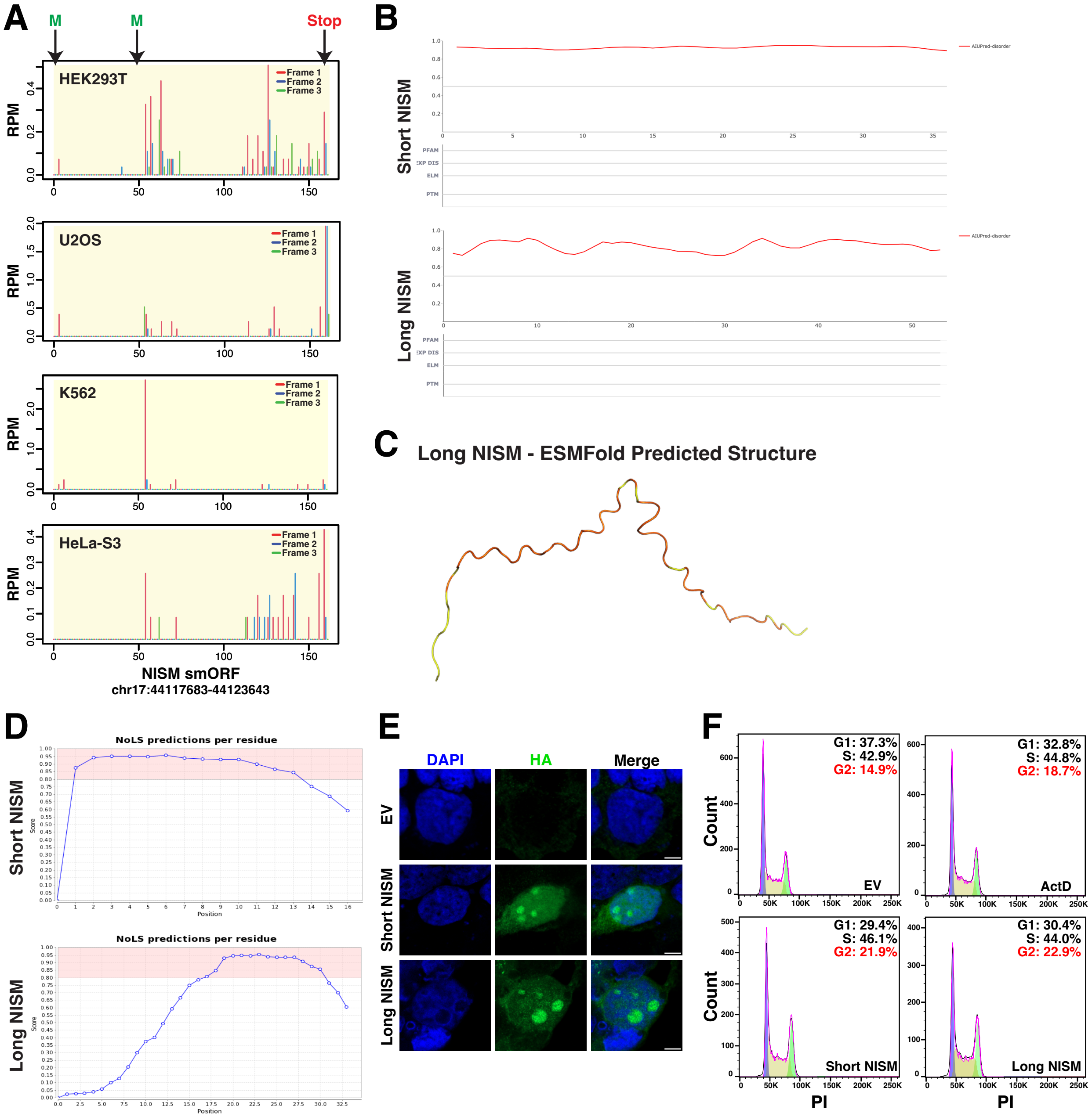


**Figure S2. Data supporting NISM's translation and characterization as an intrinsically disordered nucleolar microprotein, related to Figures 1 and 2. (A)** Ribo-seq A-site plots showing read coverage in reads per million (RPM) across all three reading frames of the NISM smORF in HEK293T, U2OS, K562, and HeLa-S3 cells. The NISM smORF is in frame 1 (red), which shows the highest coverage in each cell line. **(B)** AIUPred disordered region probability plots for each residue of short NISM (**top**) and long NISM (**bottom**). Regions scoring >0.5 are predicted to be disordered. **(C)** ESMFold predicted structure of long NISM showing no distinct structural features. **(D)** Line graphs showing the NoD nucleolar localization signal sequence probability scores for each residue of short NISM (**top**) and long NISM (**bottom**). **(E)** Immunofluorescence images of HEK293T cells transfected with empty vector (EV), short NISM-ALFA, or long NISM-ALFA for 48 h. DAPI – blue; HA – green. **(F)** Representative DNA histograms of propidium iodide stained U2OS cells treated with 10 nM ActD for 3 h, or overexpressing EV, short NISM-ALFA, or long NISM-ALFA for 72 h. All data are representative of at least two biological replicates.

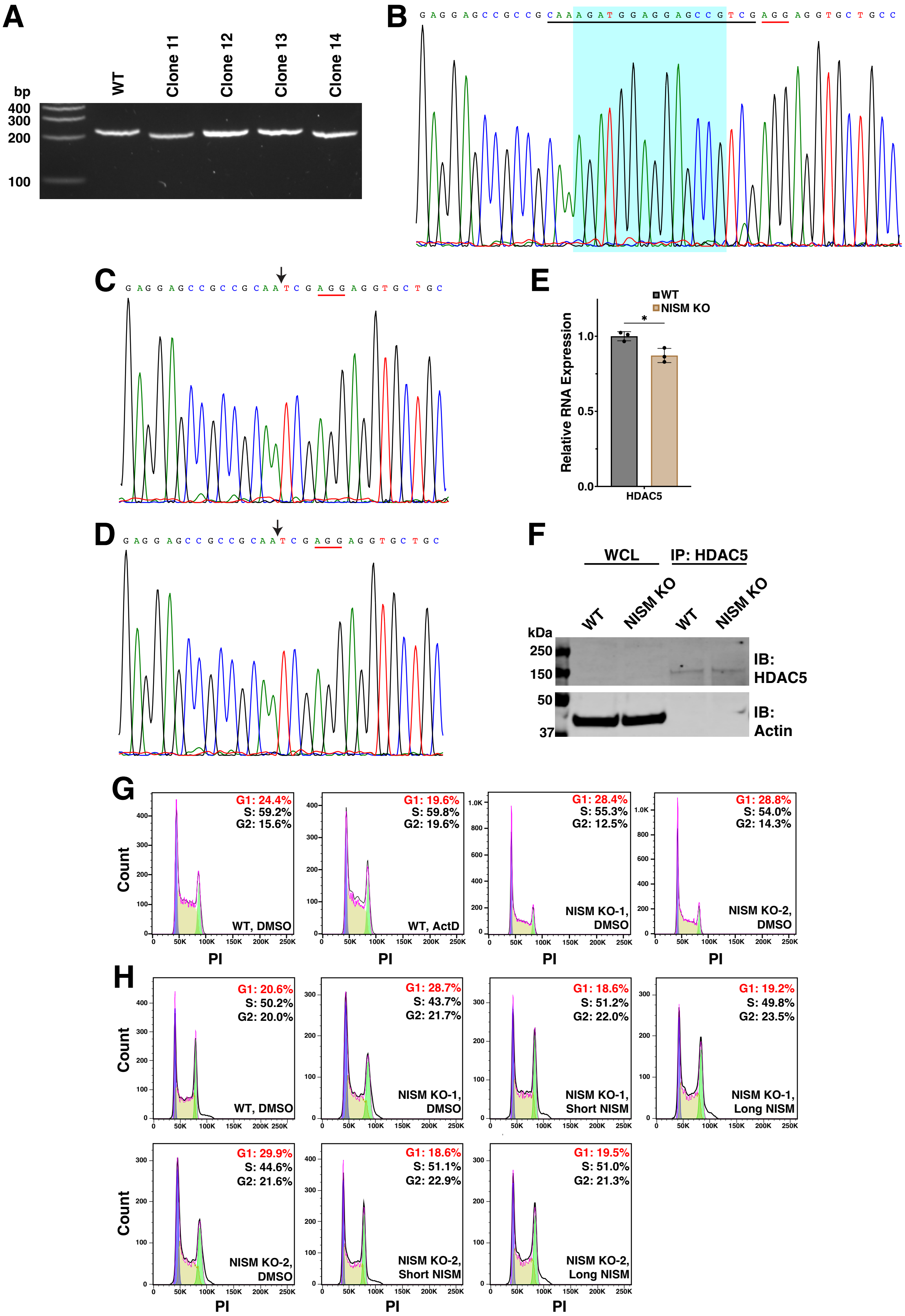

A
bp
400
300
200
100
WT
Clone 11
Clone 12
Clone 13
Clone 14
B
C
D
E
WT
NISM KO
Relative RNA Expression
HDAC5
F
WCL
IP: HDAC5
kDa
250
150
50
37
IB: HDAC5
IB: Actin
G
H
Count
PI
G1: 24.4% S: 59.2% G2: 15.6% WT, DMSO
G1: 19.6% S: 59.8% G2: 19.6% WT, ActD
G1: 28.4% S: 55.3% G2: 12.5% NISM KO-1, DMSO
G1: 28.8% S: 54.0% G2: 14.3% NISM KO-2, DMSO
G1: 20.6% S: 50.2% G2: 20.0% WT, DMSO
G1: 28.7% S: 43.7% G2: 21.7% NISM KO-1, DMSO
G1: 18.6% S: 51.2% G2: 22.0% NISM KO-1, Short NISM
G1: 19.2% S: 49.8% G2: 23.5% NISM KO-1, Long NISM
G1: 29.9% S: 44.6% G2: 21.6% NISM KO-2, DMSO
G1: 18.6% S: 51.1% G2: 22.9% NISM KO-2, Short NISM
G1: 19.5% S: 51.0% G2: 21.3% NISM KO-2, Long NISM

**Figure S3. Validation of NISM knockout in U2OS cells and cell cycle analysis raw data, related to Figure 3. (A)** Agarose gel electrophoresis showing the sizes of PCR amplicons around the NISM knockout (KO) locus for individual KO clonal cell lines. Clones 11 and 14 showed smaller single bands than the wild type (WT) control band, indicative of homozygous deletion mutants. Clone 11 is referred to as NISM KO-1 in this study and clone 14 as NISM KO-2. **(B-D)** Representative Sanger sequencing results for WT **(B)**, KO-1 **(C)**, and KO-2 **(D)**. The guide RNA targeted sequence is underlined in black, while the PAM sequence is underlined in red. The 14 bp deleted sequence in both KO-1 and KO-2 is highlighted in teal in **(B)**. The arrows in **(C)** and **(D)** depict the site where the deletion occurred in the KO clones. **(E)** Bar graphs showing the relative mRNA expression levels of HDAC5 in WT and KO-1 cells as measured by qRT-PCR. **(F)** Immunoblot showing the protein expression levels of HDAC5 in WT and KO-1 cells in HDAC5 immunoprecipitation samples. **(G-H)** Representative DNA histograms of propidium iodide stained U2OS cells associated with the bar graphs in Figures 3E, 3H, and 3I. All data are representative of at least two biological replicates.

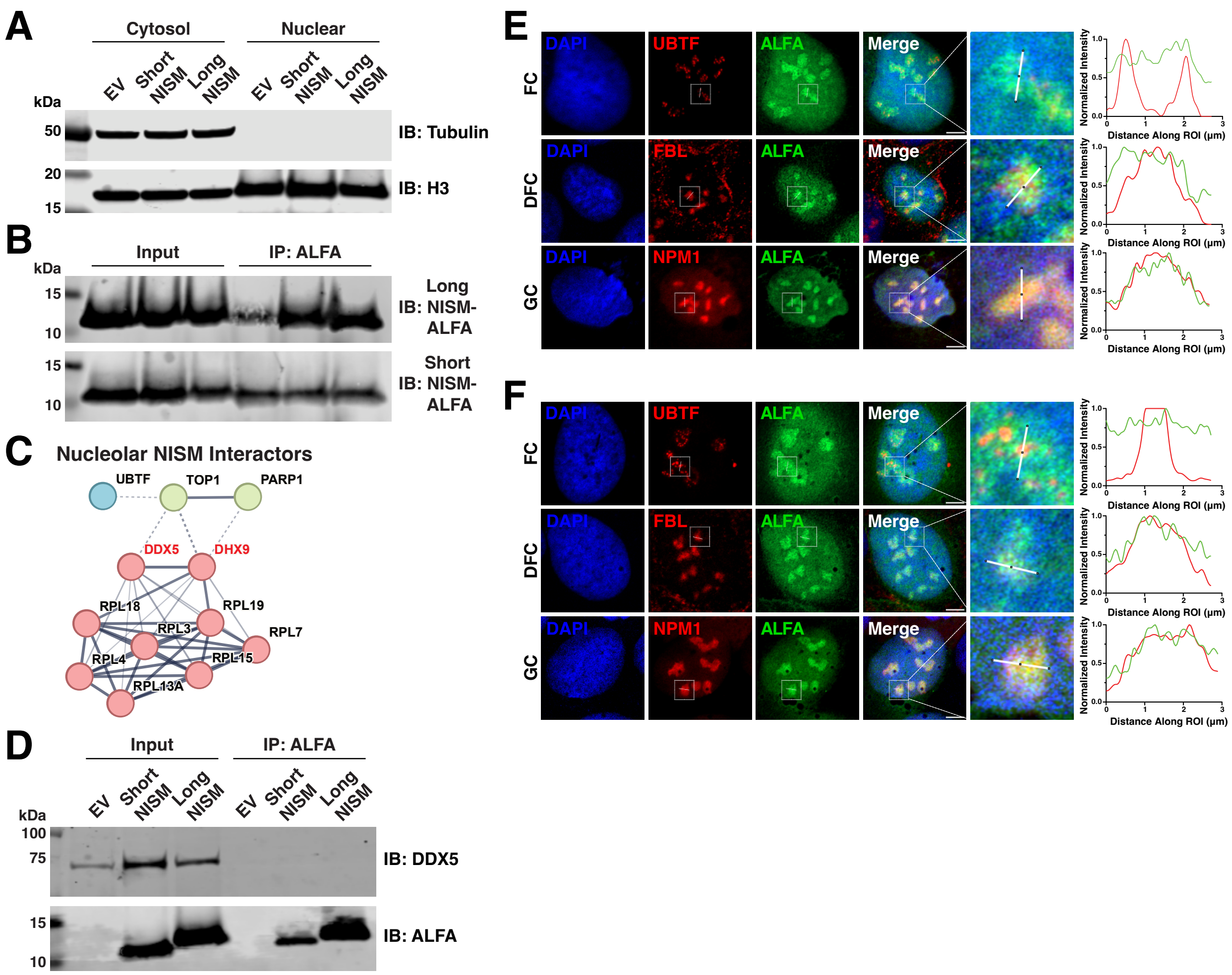


**Figure S4. Validation of NISM-ALFA nuclear IP-MS data and co-staining of NISM-ALFA with nucleolar sub-compartments, related to Figure 4. (A)** Representative immunoblots showing the enrichment of histone H3 and exclusion of tubulin from U2OS nuclear fractions used in immunoprecipitation-mass spectrometry (IP-MS) experiments. **(B)** Immunoblots showing the enrichment of short and long NISM-ALFA in IP-MS samples; n = 3. **(C)** STRING database plot showing the interactions between prey proteins pulled down with both short and long NISM-ALFA that were verified to localize to the nucleolus in the Human Protein Atlas. **(D)** Immunoblot showing no DDX5 enrichment in NISM-ALFA IP samples. **(E-F)** Representative immunofluorescence images of short NISM-ALFA **(E)** and long NISM-ALFA **(F)** co-stained with markers of different nucleolar sub-regions. UBTF is a marker for the fibrillar center (FC), FBL for the dense fibrillar component (DFC), and NPM1 for the granular component (GC). Co-localization measurements of NISM-ALFA (green) and the different sub-region markers (red) are represented by line graphs on the right. All immunoblots and images are representative of at least two biological replicates.

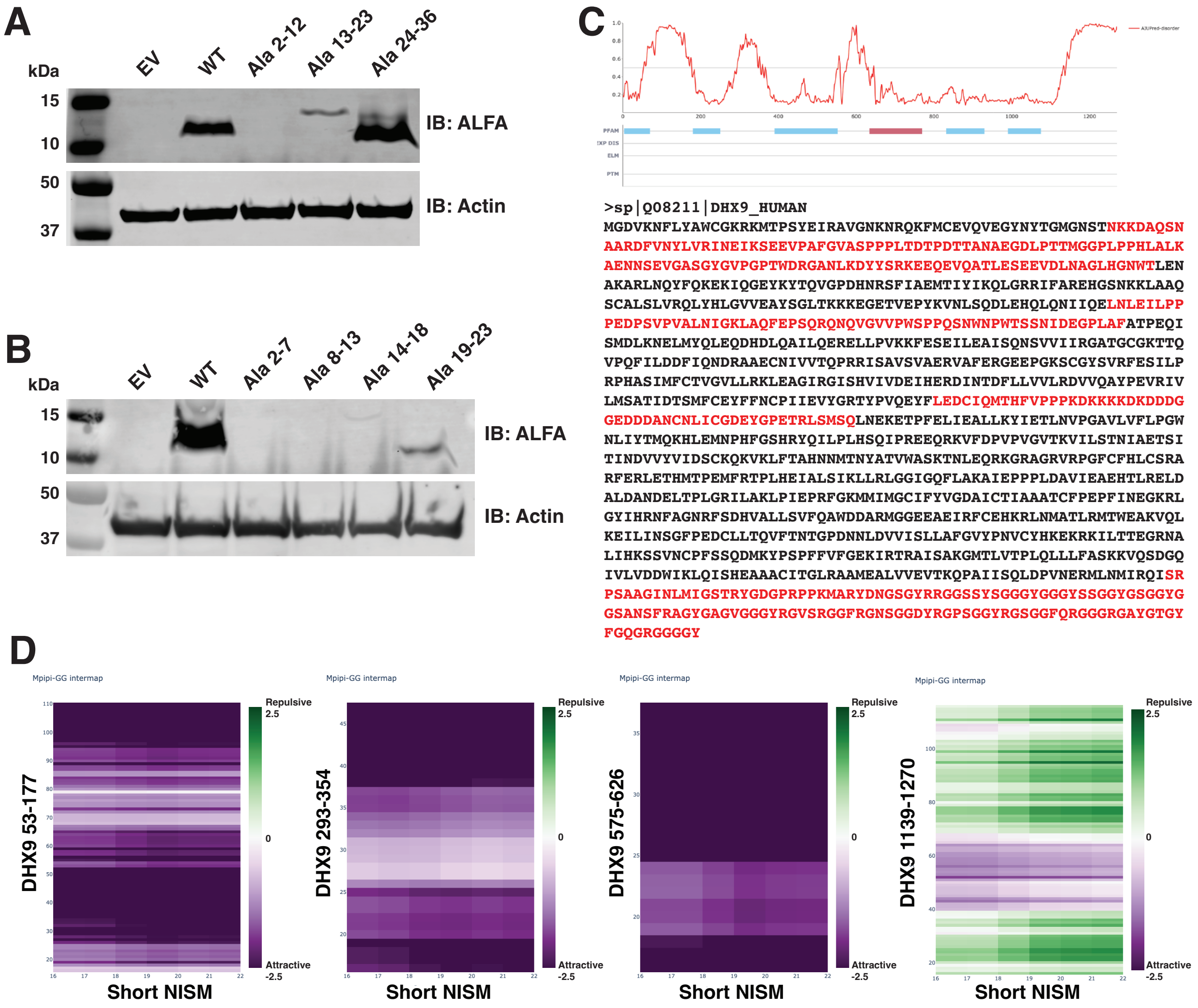


**Figure S5. Immunoblot analysis of NISM alanine mutants and computational analysis of NISM interaction with DHX9 disordered regions, related to Figure 4. (A-B)** Immunoblot analysis of NISM-ALFA alanine mutants expressed in U2OS cells. Sample names indicate which residues were mutated to alanine in each construct. **(C)** AIUPred disordered region probability plots for each residue of DHX9 (top). Regions scoring >0.5 are predicted to be disordered and are highlighted in red in the DHX9 sequence (bottom). **(D)** FINCHES Intermap plots showing the likelihood for attractive interactions (purple) and repulsive interactions (green) between short NISM and each disordered region in DHX9. All immunoblots are representative of at least two biological replicates.

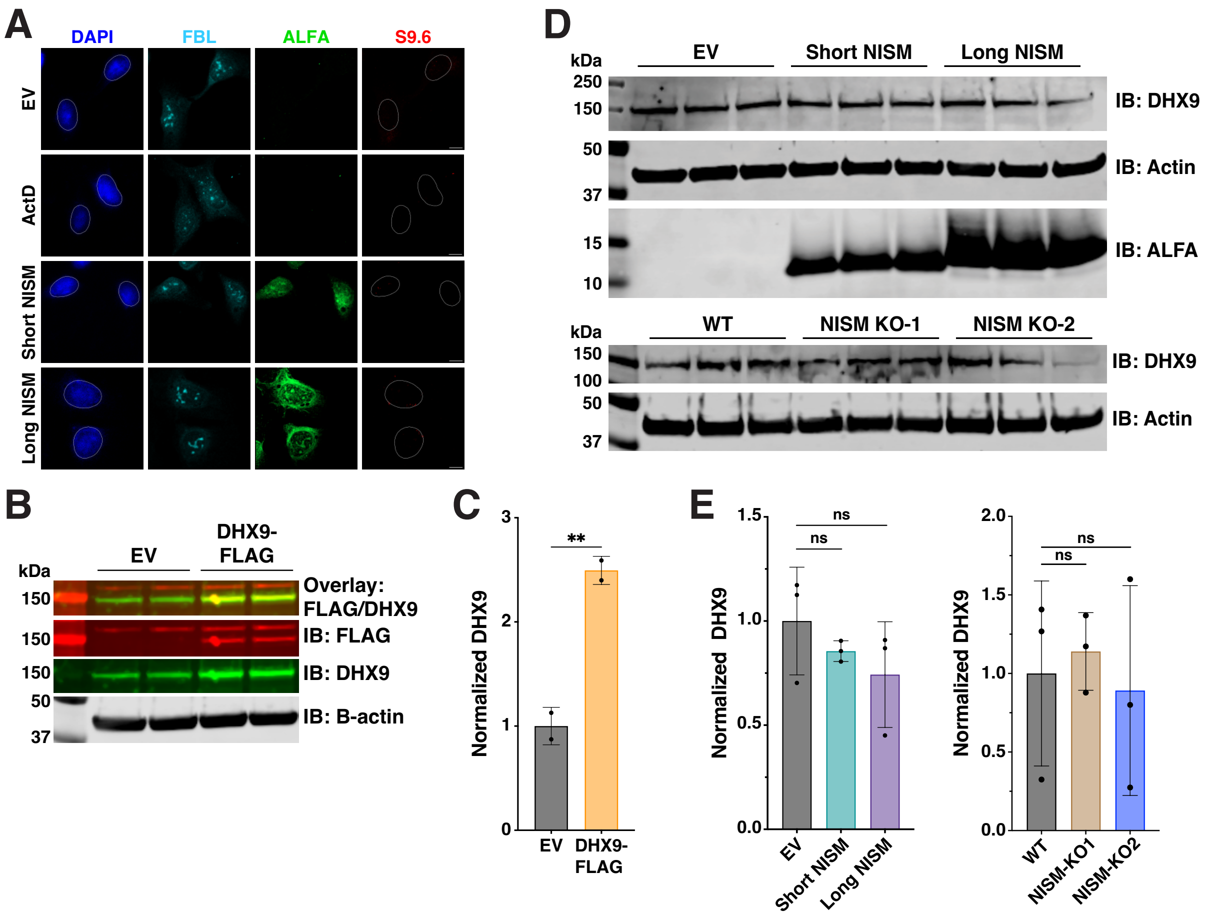


**Figure S6. RNase H control for S9.6 immunofluorescence assay and analysis of NISM perturbation on DHX9 expression, related to Figure 5. (A)** Representative immunofluorescence images of FBL (cyan), NISM-ALFA (green), and S9.6 (red) in cells treated as in Figure 6D and incubated with RNase H. RNase H abolishes S9.6 signal, demonstrating its specificity for R loops. **(B)** Immunoblot analysis of endogenous DHX9 and DHX9-FLAG in U2OS cells transfected with empty vector (EV) or DHX9-FLAG. (**C**) Quantification of DHX9 relative to Actin in (B) represented by bar plots displaying the mean ± SD. Statistical comparisons were calculated by two-tailed Student's t test; n = 2. **(D)** Immunoblot analysis of endogenous DHX9 in U2OS cells transfected with empty vector (EV), short NISM-ALFA, or long NISM-ALFA, as well as WT and NISM knockout (NISM KO) lines. (**E**) Quantification of DHX9 relative to Actin in (D) represented by bar plots displaying the mean ± SD. Statistical comparisons were calculated by two-tailed Student's t test; n = 3. ns = not significant. All data are representative of at least two biological replicates.

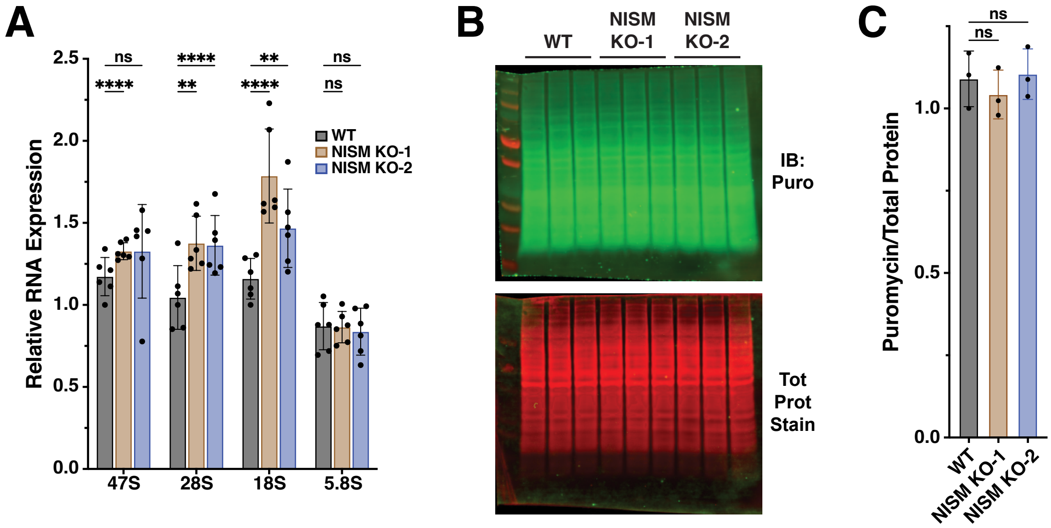


**Figure S7. NISM knockout has no effect on overall translation levels, related to Figure 6.** (**A**) Expression analysis of 47S pre-rRNA and processed 28S, 18S, and 5.8S rRNA by qRT-PCR in WT and NISM KO cells represented by bar plots displaying the mean ± SD. Statistical comparisons were calculated by two-tailed Student's t test; n = 6. **(B)** Immunoblot analysis of puromycin incorporation and total protein loading in cells as in (A). **(C)** Quantification of puromycin staining relative to total protein in (B) represented by bar plots displaying the mean ± SD. Statistical comparisons were calculated by two-tailed Student's t test; n = 3. Asterisks indicate: $^{**}p < 0.01$, $^{****}p < 0.0001$, ns = not significant. All data are representative of at least two biological replicates.

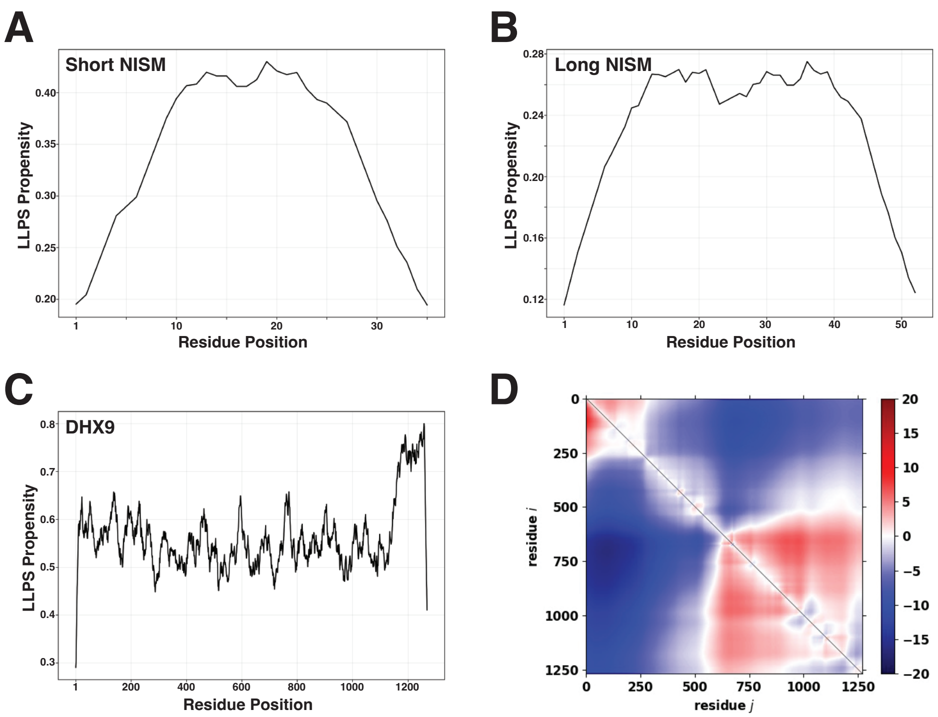


**Figure S8. Computational analyses of liquid-liquid phase separation (LLPS) propensity for NISM and DHX9, related to Figure 6. (A-C)** catGRANULE 2.0 LLPS propensity plots for short NISM (A), long NISM (B), and DHX9 (C). **(D)** Sequence Charge Decoration matrix (SCDM) comparison of DHX9 alone and DHX9 in complexation with short NISM. SCDM maps are a collection of patterning metrics that quantify sequence dependent electrostatic interaction contribution to the average distance between residue i (y axis) and residue j (x axis). Repulsive contributions are shown in red and attractive contributions are blue. The upper right triangle in the plot represents the SCDM maps of DHX9 from C-terminus to N-terminus, while the lower left triangle represents short NISM-DHX9 complexed together. For the complexed chain, only the SCDM map for DHX9 is shown for comparison with DHX9 in the absence of NISM. The lower left triangle shows darker blue regions indicating more intra-chain attraction between DHX9 residues in the presence of short NISM.
